# “Zipper” Grammar of CTD Governs the Spatial Programing of the Transcription Cycle of RNA Polymerase II

**DOI:** 10.64898/2026.01.02.697396

**Authors:** Qian Zhang, Haley A Hardtke, Yuanmin Zheng, Alan Gerber, Mukesh Kumar Venkat Ramani, Edwin Escobar, Jennifer S. Brodbelt, Ruobo Zhou, Y. Jessie Zhang

**Author notes:** Contact should be addressed to Y. Jessie Zhang.

## Abstract

RNA polymerase II transcribes all protein-coding genes in eukaryotes, with its C-terminal domain (CTD) acting as a regulatory platform throughout the transcription cycle. Despite its simple sequence, the conserved heptad repeats encode a regulatory grammar critical for transcription initiation, elongation, and termination. Using structural analysis, single-molecule imaging, and synthetic CTD-engineered Pol II constructs, we defined the minimal sequence requirements across transcriptional stages. We show that while SP motifs are essential phosphorylation sites, flanking residues tolerate variability. In contrast, the periodic positioning of tyrosine residues is indispensable for pre-initiation complex (PIC) assembly through interactions with the Mediator, which are disrupted upon Ser5 phosphorylation triggering promoter escape. This supports a “zipper” model where sequential tyrosine engagement stabilizes PIC assembly and progressive phosphorylation propels Pol II escape. Site-specific Ser2/5 phosphorylation orchestrates 3’-end processing factor recruitment. These findings redefine the functional grammar of the Pol II CTD and explain how a low-complexity sequence achieves regulatory specificity.

## INTRODUCTION

In eukaryotes, RNA polymerase II (Pol II) is the workhorse of transcription, responsible for all protein-coding mRNA as well as essential noncoding RNAs such as small nuclear RNAs (snRNAs) and small nucleolar RNAs (snoRNA) (Bartkowiak et al. 2011; Shandilya and Roberts 2012; Dignam et al. 1983). The largest subunit of Pol II contains a unique C-terminal extension absent in Pol I, Pol III, and prokaryotic and archaeal polymerases (Sklar et al. 1975; Corden et al. 1985). This unique C-terminal domain (CTD) appears to have evolved as a dedicated adaptation to facilitate Pol II transcription, enabling the enzyme to meet the high workload of protein coding gene expression. While dispensable for RNA synthesis in vitro, the CTD is essential for cell viability in vivo (M S Bartolomei et al. 1988; West and Corden 1995; Gerber and Roeder 2020), highlighting its role as a master coordinator of the transcription cycle (Eick and Geyer 2013).

The CTD is highly conserved across eukaryotes and built from tandem repeats of a heptad consensus sequence, Y_1_S_2_P_3_T_4_S_5_P_6_S_7_. Its repetitive nature suggests an origin from duplication and recombination events, favoring expansion of CTD length across evolution (Chapman et al. 2008). The number of repeats varies among species, from ∼20 in fungi to 52 repeats in vertebrates (Eick and Geyer 2013). Interestingly, only about half the full length is required for normal growth (M. S. Bartolomei et al. 1988; Thompson et al. 1993), implying that repeat multiplicity provides robustness rather than strict essentiality.

A long-standing puzzle in transcription biology is the underlying “grammar” of the CTD, such as why certain residues are strictly conserved while others are more variable and how the modification of these residues is interpreted by the transcription machinery. Among the most conserved positions of the heptad are the SP motifs where Ser2 and Ser5 undergo dynamic phosphorylation during every transcription cycle, (Ho and Shuman 1999; Zhou et al. 2012). Notably, Ser5 phosphorylation dominates at initiation (Ho and Shuman 1999), while Ser2 phosphorylation accumulates during elongation (Heidemann et al. 2013). Tyr1 is also invariant across eukaryotes: mutating all Tyr1 residues is lethal in multiple systems (Schwer and Shuman 2011; West and Corden 1995; Descostes et al. 2014; Shah et al. 2018). Recent work links Tyr1 to the CTD’s ability to undergo phase separation with transcription factors and mediator components, suggesting a role in transcriptional condensate formation (Zhang et al. 2024; Lyons et al. 2025). by contrast, Thr4 and Ser7 are more variable with mutations often tolerated (Yurko and Manley 2017; Schwer and Shuman 2011; Hintermair et al. 2012; Egloff et al. 2007).

Despite decades of study, fundamental mysteries remain unresolved. **(i)** Why do eukaryotes maintain such a long array of nearly perfect copies of the heptad consensus YSPTSPS? **(ii)** Why are phosphorylation of Ser2 and Ser5 obligatory in every transcription cycle, and how is their positional asymmetry interpreted? **(iii)** Why does nature select these seven residues, such as preserving the bulky Tyr1, precise SP motif, and the asymmetric placement of Ser2 versus Ser5 arrangement while tolerating variation at other positions? These unresolved questions point to a deeper logic in CTD sequence design beyond its role as a generic “landing pad” for transcription-associated factors.

Recent advances in cryo-electron microscopy now allow direct visualization of Pol II-Mediator interactions at near atomic detail. Building on these structural insights, we systematically engineered Pol II variants with synthetic CTDs containing defined alterations in heptad composition and spacing. Our analysis reveals that approximately one-third of the CTD length is sufficient to mediate preinitiation complex (PIC) assembly in human cells. Within this segment, periodic Tyr1 residues engage the Mediator through hydrophobic interactions, anchoring the CTD to stabilize the PIC. Extensive phosphorylation of adjacent SP motifs disrupts these interactions, promoting Pol II escape from promoter. Strikingly, swapping Ser2 and Ser5 did not impair initiation or pause release, but selectively compromised elongation, underscoring that Tyr periodicity, rather than Ser positioning, is the dominant determinant of PIC stability. Overall, these findings support a “zipper model” in which unphosphorylated CTD Tyr residues sequentially engage the Mediator, requiring at least one-third of the native CTD to remain unphosphorylated. Phosphorylation of SP motifs then pries open these interactions, driving promoter clearance and transcriptional progression. This model explains how a low-complexity sequence encodes a positional and dynamic code, resolving long-standing questions about why the CTD’s repetitive grammar is both conserved and essential for eukaryotic transcription.

## RESULTS

### Structural modeling of the minimal CTD length for human PIC assembly

One notable feature of the Pol II CTD is that the repeat number varies across eukaryotes, yet not all repeats are necessary (Eick and Geyer 2013). With 26 repeats in Pol II, *S. cerevisae* only requires ∼9-11 CTD repeats to support growth (Ling et al. 2024; West and Corden 1995; Nonet and Young 1989), while the precise number of CTD repeats required to support human cell growth has been debated (M S Bartolomei et al. 1988). To address this, we first analyzed recently published PIC complex structures to better understand the role of the CTD in PIC assembly. In most Pol II structures, the CTD remains invisible due to its inherent flexibility. However, advancements in cryo-EM have enabled the capture of Pol II at multiple transcriptional stages (Abril-Garrido et al. 2023; Chen et al. 2022; Schilbach et al. 2023; Ehara et al. 2019; 2022; Farnung et al. 2021; 2022; Filipovski et al. 2022; Chen et al. 2023). Importantly, two PIC structures (yeast at 3.6 Å and human at 4.2 Å resolution) resolved portions of the CTD bound to Mediator (Schilbach et al. 2023; Chen et al. 2022). The location and binding mode of the CTD are strikingly conserved: fragments of CTD heptads wrap around the interface between the Mediator head and middle modules, threading through MED6, MED8, and MED17 (head) and MED4, MED7, MED14, and MED31 (middle) (Figure 1A, SI 1A-1B). In the absence of CTD, the Mediator head-middle interface is tenuous, with an interface of 1,292.4 Å^2^ in the human PIC, creating obvious gaps (Figure 1A, SI 1C). When the CTD fragments are present, the heptad repeats bridge this gap, forming extensive interactions with both modules (Figure 1A, SI 1A-1B). Our model indicates that the CTD-Mediator interface spans 3528.5 Å^2^ with 1585.2 Å^2^ involving the Mediator head and 1943.3 Å^2^ involving the Mediator middle (Figure SI 1D). Thus, the CTD acts as a structural “glue”, stabilizing head-middle interactions of the Mediator to promote PIC assembly (Figure 1A).

**Figure 1:**
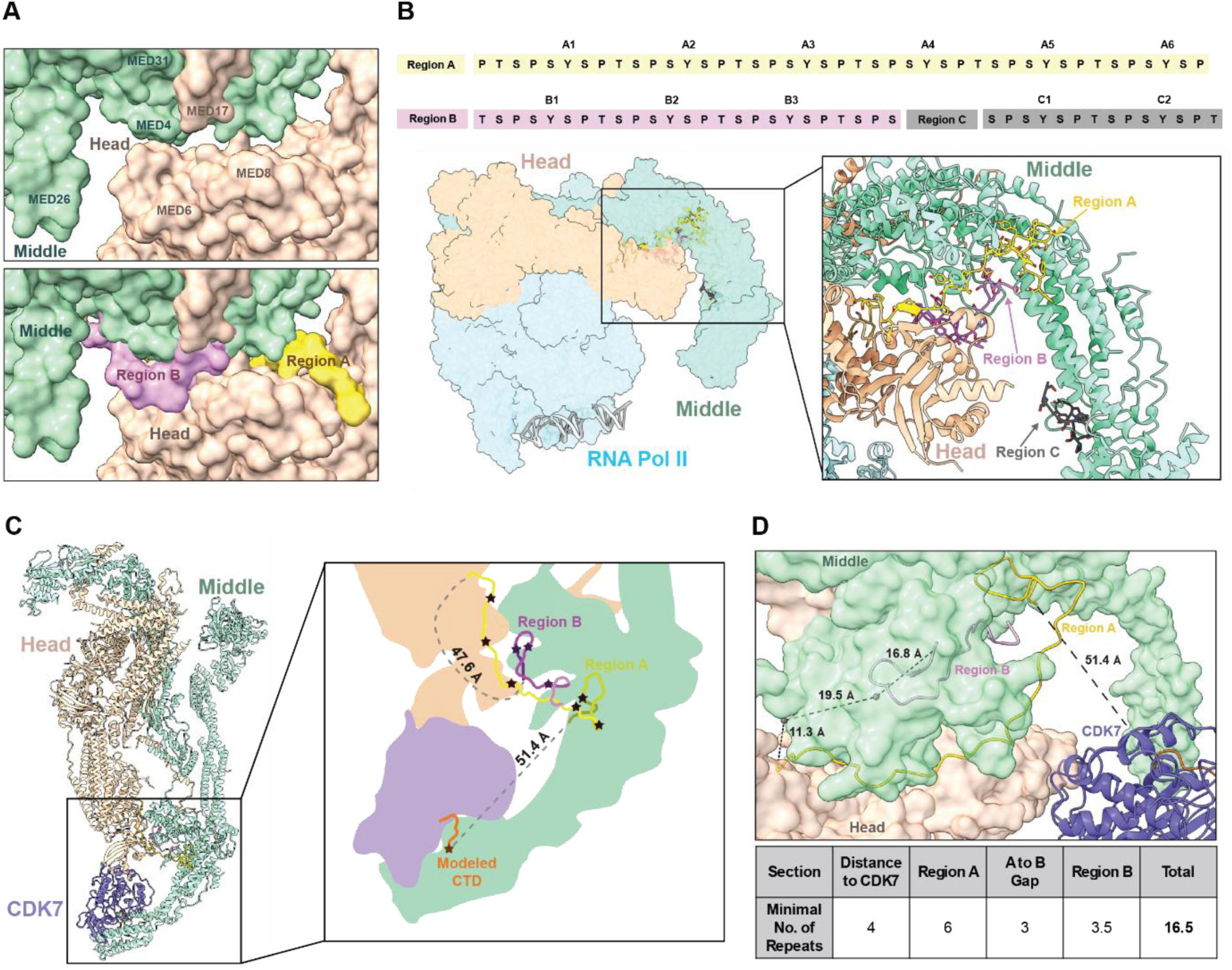
Structural modeling of minimal length of CTD for human PIC assembly. (A) Model of the human Mediator bound to consensus CTD repeats. The top figure shows the surface contacts between the Mediator Head and Middle regions without the CTD, and the bottom figure shows the same region with the CTD present. (B) Structure of yeast Pol II in complex with the yeast Mediator Head and Middle regions (PDB Code: 8CEO). A close-up view of the interaction between the yeast Mediator and the Pol II CTD is shown as a cartoon. The sequences of the three visible stretches of the CTD (named regions A, B, and C) are listed at the top of the figure. (C) Top view of the human Mediator-bound to consensus CTD repeats in complex with CDK7. The yeast PIC (PDB Code: 8CEO) was superimposed on the human PIC (PDB code: 8GXS) to position the repeats in the human structure. A model of CDK7 bound to the Pol II CTD was superimposed onto the apo CDK7 within the human PIC structure. The distances between regions A and B as well as region A and CDK7 are shown in the cartoon schematic to the right. The stars in the cartoon schematic indicate the location of tyrosines in the CTD chains. (D) Model of the human Mediator bound to the consensus CTD repeats. To measure the distance between regions A and B we placed markers along the surface of the Mediator Head to estimate the distance from region A around the Mediator Head to region B. A model of CDK7 bound to a CTD peptide was superimposed onto CDK7 in the human PIC structure (PDB code: 8GXS), and the distance from region A to the CDK7 active site was also calculated.

To define the minimal CTD length, we analyzed the known yeast PIC structure, which provides the best resolution (Schilbach et al. 2023) (Figure 1B). Since the linker region which connects the core of Pol II to the CTD is completely disordered, it is impossible to determine the precise repeat number or orientation of the heptad. We thus named the fragments captured within the structure regions A, B, and C, and numbered the tyrosines within each fragment (Figure 1B). Region A, the main interaction site, contains ∼5 full and two partial repeats and bridges the head-middle subunits. Region B locates on the opposing side of the “neck” between the two modules and comprises ∼3.5 repeats (Figure 1B). Region C, visible only in the yeast PIC and contacting poorly conserved Mediator middle subunits, was excluded from further analyses due to uncertain relevance. In addition to Mediator binding, the CTD must extend to the kinase module, as CDK7 phosphorylation is required for transcription activation (Lu et al. 1992). Using the human PIC structure with an attached kinase module for reference (Chen et al. 2022), we positioned an activated CDK7, by analogy with CDK2 (Wood and Endicott 2018; Ramani et al. 2020), relative to the Mediator position in our model. Modeling a heptad into the CDK7 active site allowed us to measure the distance from CDK7 to Region A, yielding 51.4 Å (Figure 1C).

Given that one heptad spans ∼8–24 Å depending on conformation, we estimated repeat requirements for each segment. Region A and B account for ∼6 and ∼3.5 repeats, respectively. The gap between A and B is 47.6 Å (∼3 repeats). The distance from Region A to the CDK7 active site requires ∼3 repeats, to which we added an additional repeat to ensure accessibility. These results suggest that a minimal CTD length of ∼16–17 repeats are sufficient to support human PIC assembly and initiation of transcription (Figure 1C-1D).

### Experimental testing of a minimal CTD length for cell survival

To experimentally interrogate our structural prediction for the minimal CTD length, we performed cell growth assays with RPB1 constructs harboring different numbers of CTD repeats. Previous studies in human cells yield inconsistent results, largely due to the incomplete removal of endogenous RPB1 (McCracken et al. 1997; Sawicka et al. 2021; Meininghaus et al. 2000; Meininghaus and Eick 1999). To overcome this limitation, we employed an inducible auxin-degron system to degrade endogenous RPB1-AID while transiently overexpressing RPB1 variants with defined CTD lengths (Figure SI 2A). In HEK293 cells, a stable cell line was generated expressing full-length RPB1 fused at the C terminus with a minimal auxin-sensitive degron while endogenous RPB1 was removed using CRISPR-Cas9. In this RPB1 degron background, robust auxin-dependent degradation of RPB1-AID occurred upon doxycycline (Dox)-induced expression of the ubiquitin ligase OsTIR1 for 12 hours, followed by a 2 hour auxin treatment (Gerber et al. 2020). To determine the minimal CTD length to support cell viability, we constructed eight RPB1 variants containing 0, 3, 9, 15, 17, 20, 26, or 52 CTD heptad repeats. Each construct was transiently overexpressed in HEK293 RPB1 degron cells for 48 hours, followed by auxin treatment to degrade RPB1-AID (Gerber et al. 2020). Cells were seeded at equal densities at time 0, and viable cell numbers were measured 96 hours later. The 0CTD construct exhibited a lethal phenotype (Figure 2A). Similarly, 3CTD, 9CTD, and 15CTD variants supported minimal growth, indistinguishable from 0CTD. Strikingly, the 17CTD variant resulted in a statistically significant increase in cell number compared to the shorter CTD variants, with cell counts similar to those of the 20CTD and 26CTD constructs (Figure 2A).

**Figure 2:**
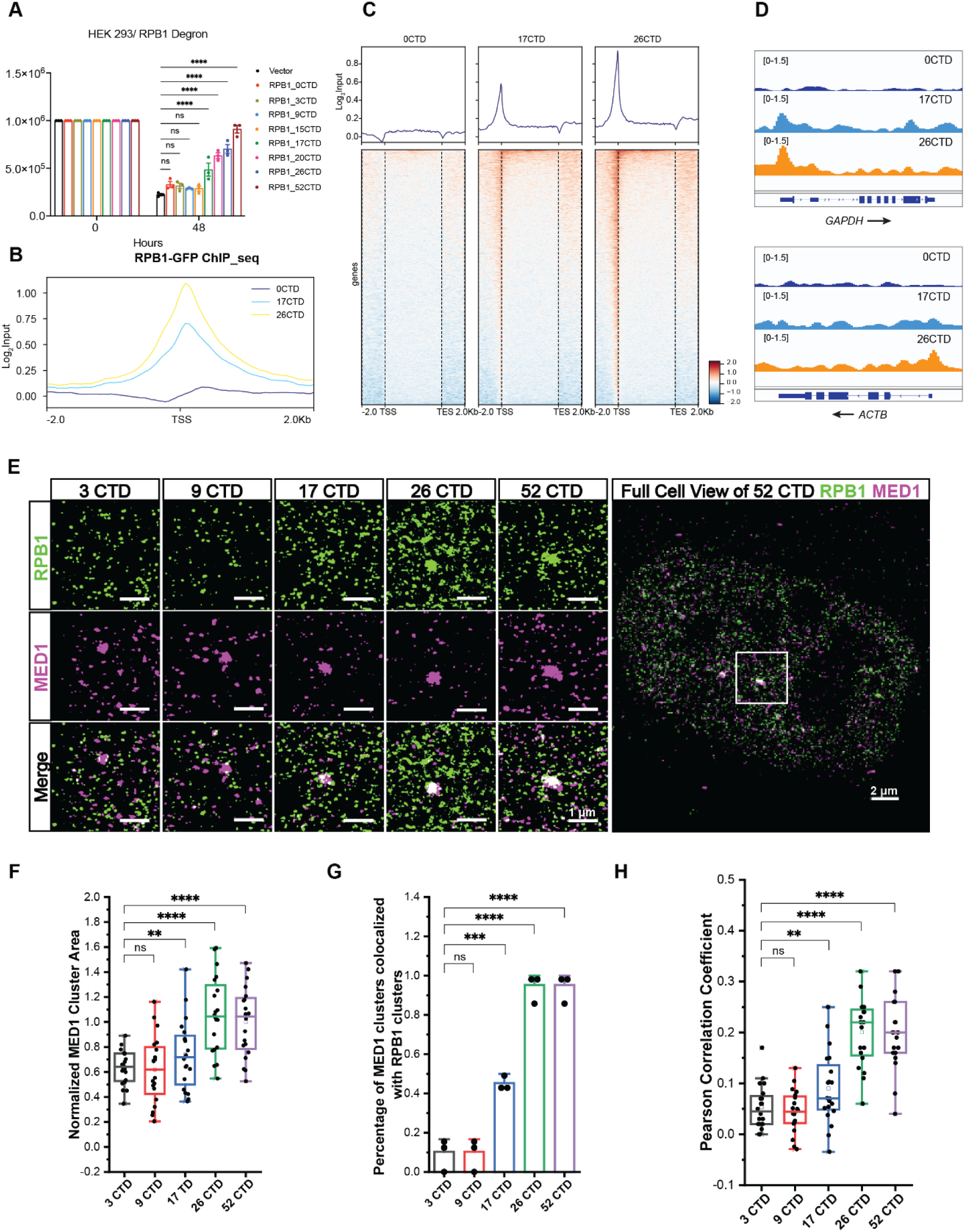
The minimal length of CTD for cell survival and PIC assembly in cells. (A) Cell density in transfected RPB1 with different CTD length upon auxin administration in HEK 293 RPB1 degron cells. n = 3 independent experiments (mean ± SEM); ****p < 0.0001, comparison was performed using two-way ANOVA with Tukey correction. (B) Heatmaps from ChIP-seq analysis of RPB1 0CTD and 17 CTD constructs. Two biological replicates (n=2). (C) Merged metagene plot of RPB1 ChIP-seq data. (D) Browser tracks of RPB1 ChIP-seq data at GAPDH and ACTB. (E) Dual-color STORM images of RPB1 (Green) and MED1 (Magenta) clusters. Scale bars: 1 μm. U2OS cells were transfected with YFP-tagged RPB1 constructs containing 3, 9, 17, 26, or 52 CTD heptad repeats. (F) Normalized MED1 cluster area for the five RPB1 constructs. Boxplots show the mean, median and boundaries (first and third quartile); the whiskers denote the minimum and maximum excluding outliners. For each cell, the largest MED1 cluster was measured and normalized to the 52CTD group. Each data point represents a single cell. Comparison was performed using one-way ANOVA with Tukey correction. (G) Percentages of the MED1 clusters that show colocalization with RPB1 clusters for the five RPB1 constructs. For analysis, 2 μm × 2 μm regions containing MED1 clusters were cropped, and the largest RPB1 and MED1 clusters were compared. Clusters were defined as colocalized when the overlapping area was > 0. Data are presented as mean ± SEM. (n = 3 biological replicates per condition). Comparison was performed using one-way ANOVA with Tukey correction. (H) Pearson correlation coefficients between MED1 and RPB1 for the five RPB1 constructs. The largest MED1 and RPB1 puncta within each cropped region were used for calculation. Each data point represents a single cell. Boxplots as in (F). Comparison was performed using one-way ANOVA with Tukey correction.

To further validate these findings, we employed an independent α-amanitin–based complementation system. RPB1 variants of varying CTD lengths were engineered in the background of the α-amanitin–resistant N792D RPB1 mutant (Nguyen et al. 1996). Following 24 hours of expression, α-amanitin was added to eliminate endogenous Pol II, and cell viability was assessed 72 hours later. Results closely mirrored those from the degron system (Figure SI 2B). Together, both approaches consistently demonstrate that ∼17 CTD heptad repeats are sufficient to support human cell survival.

Based on this, we performed ChIP-seq analysis to identify the genomic locations of Pol II variants harboring either 0 or 17 CTD heptad repeats. Each Pol II mutant was immunoprecipitated using an anti-GFP antibody to specifically map the distribution of Pol II with different CTD lengths. Genome-wide profiles revealed that RPB1 with 17 CTD repeats was enriched at promoter regions, comparable to RPB1 with 26 CTD repeats. In contrast, the 0CTD mutant completely lost chromatin binding across the genome, most notably at transcription start sites (TSS) (Figure 2B-2D), indicating that the CTD is essential for PIC assembly and that approximately 17 CTD heptad repeats are sufficient to support PIC assembly in human cells.

### Super-resolution imaging of PIC assembly

To further investigate how CTD length influences the PIC assembly, we used two-color stochastic optical reconstruction microscopy (STORM) (Sigal et al. 2018; Zhou et al. 2019) to examine the nanoscale spatial relationship between endogenous Mediator (MED1) and transiently expressed exogenous RPB1 constructs containing 3, 9, 17, 26, or 52 CTD heptad repeats in U2OS cells. Endogenous MED1 was immunostained with a specific anti-MED1 antibody (Zhang et al. 2024), while exogenous YFP-tagged RPB1 was selectively detected using an anti-YFP antibody (Zhou et al. 2022), ensuring that only the exogenous, rather than endogenous, RPB1 was visualized (Figure 2E, Figure SI 2C). Although MED1 clusters were detectable under all five CTD length conditions due to the presence of endogenous RPB1, we observed a progressive reduction in average MED1 cluster area as the CTD length decreased, with a marked decline when the number of heptad repeats dropped to 17 or fewer (Figure 2F). These observations suggest that exogenous RPB1 competes with endogenous RPB1 for PIC formation, and that shorter CTDs (≤17 repeats) are less efficient in recruiting MED1.

To quantitatively assess RPB1-MED1 colocalization (i.e., the extent of exogenous RPB1 recruitment to the PIC) for the five CTD length conditions, we calculated three metrics from the STORM images: 1) the percentage of MED1 clusters overlapping with RPB1 clusters; 2) the Pearson correlation coefficient between the two color-channels; and 3) the fraction of MED1 cluster area overlapping with RPB1 signal. All three metrics revealed a progressive decline in colocalization as the number of CTD repeats decreased (Figure 2G and 2H, SI 2D). Notably, colocalization dropped significantly at 17 repeats and approached zero in cells expressing the 9CTD or 3CTD constructs.

Taken together, our structural analysis of the Pol II PIC and our experimental results reveal 17 repeats as the critical number of repeats required to support PIC assembly, CDK7-mediated CTD phosphorylation, and cell growth. The CTD of Pol II ties together the head and middle modules of the Mediator, stabilizing the complex formation during PIC assembly. Thus, omission of the CTD is lethal for human cells, likely due to the loss of stable PIC formation.

### CTD drives transition from assembled PIC to promoter clearance

After PIC assembly, the CTD becomes processively phosphorylated at Ser5 as transcription initiated, although it remains unclear from which end this phosphorylation begins (Søgaard and Svejstrup 2007; Jeronimo and Robert 2014; Wong et al. 2014). To dissect how the CTD transitions from an unphosphorylated to a phosphorylated state, we engineered RPB1 constructs in which one portion of the CTD remained unphosphorylated while the other half mimicked phosphorylation at Ser5 using a glutamate. Given that the 52 heptad repeats of the human CTD contain more conserved consensus sequences in the proximal region and greater variability in the distal region, we designed the phospho-mimicking S5E substitution at the proximal or distal half of the human CTD (Figure 3A).

**Figure 3:**
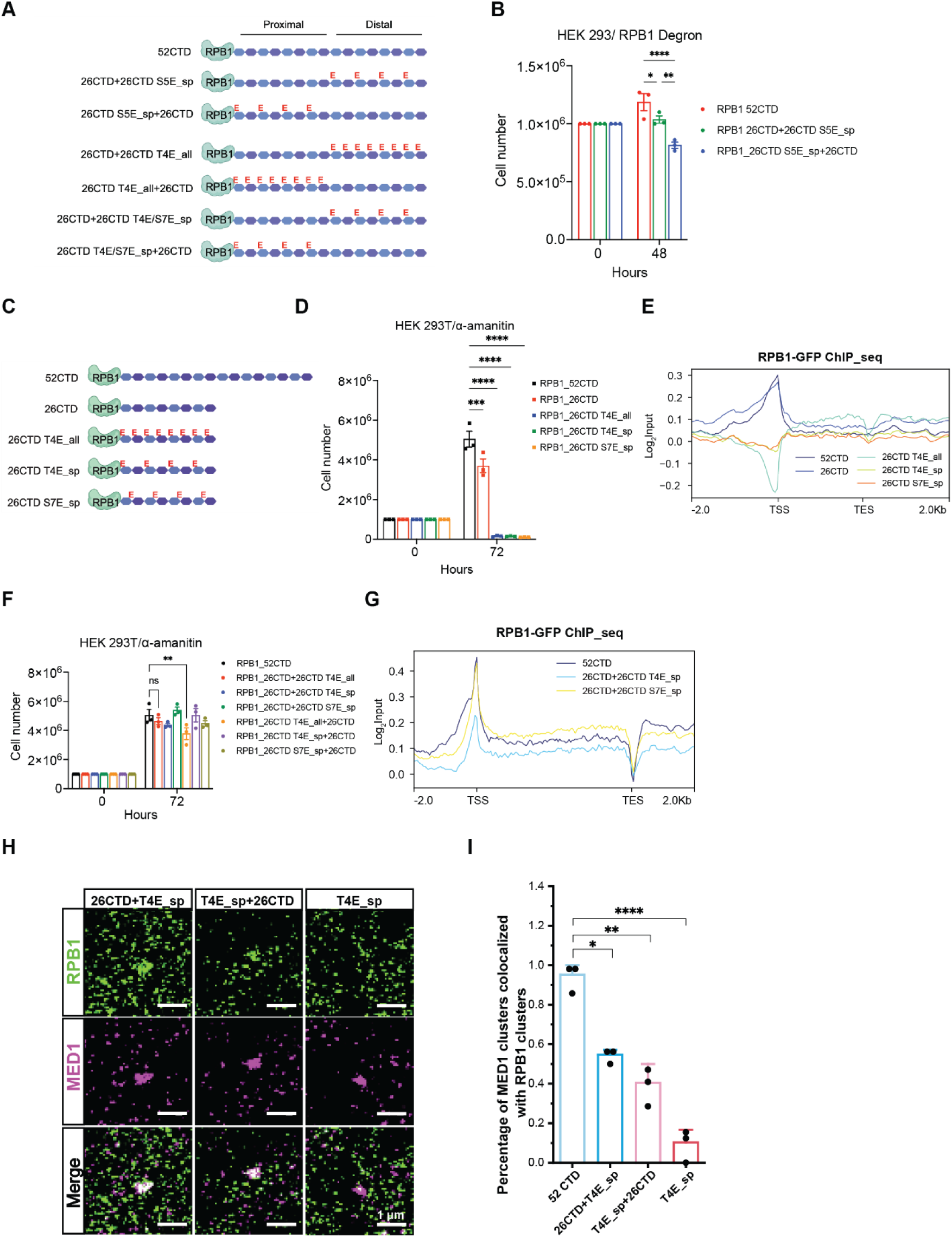
Phosphorylation of CTD disassembles PIC and facilitates initiation. (A) Schematic of RPB1 52CTD (WT) and hybrid CTD variants. (B) Cell density after transfection of RPB1 with 52CTD (WT) or different 26CTD S5E_sp hybrid constructs in HEK 293 RPB1 degron cells upon auxin administration. n = 3 independent experiments (mean ± SEM); ****p < 0.0001, **p < 0.01, *p < 0.05, comparison was performed using two-way ANOVA with Tukey correction. (C) Schematic of RPB1 52CTD (WT) and multiple CTD variants. (D) Cell density after transfection of RPB1 with different CTD constructs in HEK293T cells after α-amanitin administration. n = 3 independent experiments (mean ± SEM); ****p < 0.0001, ***p < 0.001, comparison was performed using two-way ANOVA with Tukey correction. (E) Merged metagene plot of RPB1 ChIP-seq data with different CTD mutant constructs. (F) Cell density after transfection of RPB1 with different hybrid CTD constructs in HEK293T cells after α-amanitin administration. n = 3 independent experiments (mean ± SEM); **p < 0.01, comparison was performed using two-way ANOVA with Tukey correction. (G) Merged metagene plot of RPB1 ChIP-seq data with different hybrid CTD constructs. (H) Dual-color STORM images of RPB1 (Green) and MED1 (Magenta) clusters. Scale bars, 1 μm. U2OS cells were transfected with YFP_RPB1 26CTD+26CTD T4E_sp, 26CTD T4E_sp+26CTD, and 26CTD TE4_sp. (I) Percentages of the MED1 clusters that show the colocalization between MED1 and RPB1 for the three constructs indicated in (H). Data are presented as mean ± SEM (n = 3 biological replicates per condition). Comparison was performed using one-way ANOVA with Tukey correction.

We expressed these RPB1 variants in RPB1 degron cells, suppressing endogenous Pol II to test whether they could support cell growth. Both mutants were viable but displayed slower growth than wild-type RPB1, with a slightly greater defect when the proximal half contained the phospho-mimicking substitution (Figure 3B). These results indicate that even when half of the human CTD is phosphorylated, the remaining unphosphorylated half can still support transcription, regardless of which end the CTD phosphorylation starts (Figure 3B).

Previous genetic studies have shown that substituting almost any CTD heptad residue with a phospho-mimicking amino acid in each repeat, thereby simulating a fully phosphorylated CTD, is lethal both in yeast and human cells (Schwer and Shuman 2011, 1; Harlen et al. 2016; Kecman et al. 2018; Zhang et al. 2012). Given that Ser2 and Ser5 phosphorylation are essential for transcription, their dominant lethality is not hard to understand, however, it is surprising that phospho-mimicking substitutions of residues such as Ser7 or Thr4, which are poorly conserved across species and generally tolerate most mutations, also exhibit lethal phenotypes (Schwer and Shuman 2011; Hintermair et al. 2012; Egloff et al. 2007). To pinpoint which step of transcription requires CTD dephosphorylation, we generated RPB1 variants in which Thr4 or Ser7 were systematically replaced with glutamate, either in every repeat or in alternating repeats to model different levels of phosphorylation (Figure 3C).

We expressed these RPB1 variants in an α-amanitin-resistant system, suppressing endogenous Pol II to test whether they could support cell growth. In all cases, cells expressing phospho-mimetic CTDs failed to survive (Figure 3D), demonstrating that the mutants were universally non-functional. To investigate the basis of this defect, we performed ChIP-seq to map the genomic locations of these variants RPB1. The phospho-mimetic Pol II mutants were absent at gene promoters (Figure 3E). To understand why CTD phospho-mimetic variants cannot bind to promoters, we conducted structural modeling of the PIC when such mutation occurs. Structural modeling revealed that phosphorylation at any position within CTD regions A or B across consecutive repeats generates severe steric clashes that disrupt the Mediator head-middle module interface (Figure SI 3A). Even phosphorylation in alternating repeats destabilizes the interactions as the interface is dominated by hydrophobic and non-polar interactions. Thus, it is not the position of phosphorylation, rather the consistent negative charge that prevents PIC assembly.

To test if consistent negative charge rather than the specific position of the phosphorylation causes the lethal phenotype, we generated an RPB1 construct in which the first half of the CTD was unmodified while the other half carried glutamate substitution at Thr4 (26CTD+26CTDT4E_all) (Figure 3A). Since phosphorylation during initiation is not necessarily uniform, we also created hybrid constructs in which only every other repeat in the second half was mutated to mimic partial phosphorylation (26CTD+26CTDT4E_sp, 26CTD+26CTDS7E_sp). To avoid positional bias, if the phosphorylation occurs at the proximal or distal half, we also reversed the order of the mutant and wild-type halves (26CTDT4E_all+ 26CTD) (Figure 3A). Using an α-amanitin–resistant system, we tested whether these hybrid CTDs could sustain cell growth in the absence of endogenous Pol II. Remarkably, all the hybrid constructs supported viability with only modest or negligible growth defects compared to cells expressing the wild-type 52CTD (Figure 3F). ChIP-seq analysis further revealed that, unlike the complete loss of promoter occupancy seen in fully phospho-mimicking mutants, the hybrids restore Pol II binding the at transcription start site (TSS) (Figure 3G, SI 3B). These findings indicate that unphosphorylated CTD repeats are specifically required for promoter recruitment and that this function can be fulfilled by a defined CTD segment within the CTD tail.

### STORM imaging of the RPB1 mutants mimicking the transition from initiation to elongation

To determine whether the hybrid or phospho-mimicking CTDs support PIC assembly, we performed two-color STORM imaging in U2OS cells transiently expressing YFP-tagged RPB1 hybrid CTD constructs. In contrast to a variant carrying phospho-mimetic substitutions across the entire 26CTD (26CTD T4E_sp) which exhibited no MED1 clustering, the hybrid constructs (26CTD+26CTDT4E_sp and 26CTDT4E_sp+26CTD) exhibited clear MED1 co-localization (Figure 3H, SI 3B). Although the average MED1 cluster area was smaller than with the 52CTD, it remained significantly larger than with the 26CTD T4E_sp mutant (Figure 3I). These results suggest that hybrid CTDs, like the 17CTD construct, support cell viability despite a moderate deficiency in PIC assembly. We next quantified RPB1–MED1 colocalization using three STORM-based metrics defined earlier. All analyses revealed that the hybrid constructs displayed intermediate levels of colocalization, indicating partial recruitment of RPB1 to MED1 clusters (Figure SI 3C-3F). Interestingly, the 26CTD+26CTD T4E_sp construct showed slightly higher colocalization than its reverse-order counterpart, suggesting a positional effect of the unphosphorylated segment on PIC assembly efficiency. By contrast, the fully phospho-mimetic 26CTDT4E_sp mutant exhibited minimal colocalization, similar to short CTD constructs (3CTD, 9CTD), confirming that extensive phosphorylation severely impairs PIC recruitment.

Together, these results recapitulate the transition from PIC assembly to promoter clearance. PIC assembly requires an unphosphorylated stretch of CTD heptad repeats long enough to engage the Mediator head and middle modules. As transcription initiates, progressive phosphorylation gradually proceeds, introducing steric and chemical clashes that destabilize Mediator–CTD contacts. This phosphorylation eventually “pries open” the PIC, ejecting Pol II. This model provides a structural mechanism for Pol II escape into productive elongation.

### Hydrophobic interaction by Tyr1 is crucial for PIC assembly

In the PIC structures, Tyr1 threads snugly through the interface between the Mediator head and middle modules, where periodically placed hydrophobic pockets accommodate Tyr1 side chains roughly one heptad apart (Figure 4A) (Schilbach et al. 2023). These interactions are primarily hydrophobic, with additional hydrophilic contributions from Tyr1 hydroxyl groups. The bulkier Tyr1 residues anchor the CTD heptad repeats into hydrophobic Mediator pockets while each hydrophobic pocket is about seven residues apart on the surface of the Mediator (Figure 4A). In region A, the first Tyr1 (A1) inserts into a deep pocket formed by MED8 and MED17. Its aromatic ring is sandwiched by Leu109/113 of MED8 and Leu220 of MED17, while its hydroxyl group hydrogen bonds with Asp223 of MED17 (Figure 4B, left). A second Tyr1 (A2) from the adjacent heptad anchors into a neighboring cleft formed by MED6 and MED8, stacking against Tyr161 of MED8, which itself stacks with Tyr154 (Figure 4B, right). In region B, located on the opposite side of the head–middle neck, Tyr1 (B1) fits into a MED31 pocket supported by Val122 (MED31), Trp182 and Pro177 (MED4), with its hydroxyl group positioned near Asn119 of MED31 (Figure SI 4A, left). Tyr1 (B2) interacts with MED6, potentially forming a hydrogen bond with Asp124, while Tyr1 (B3) anchors in a MED4 pocket lined by Phe180, Pro167, and Val175, with Lys162 orienting the hydroxyl group (Figure SI 4A, middle and right).

**Figure 4:**
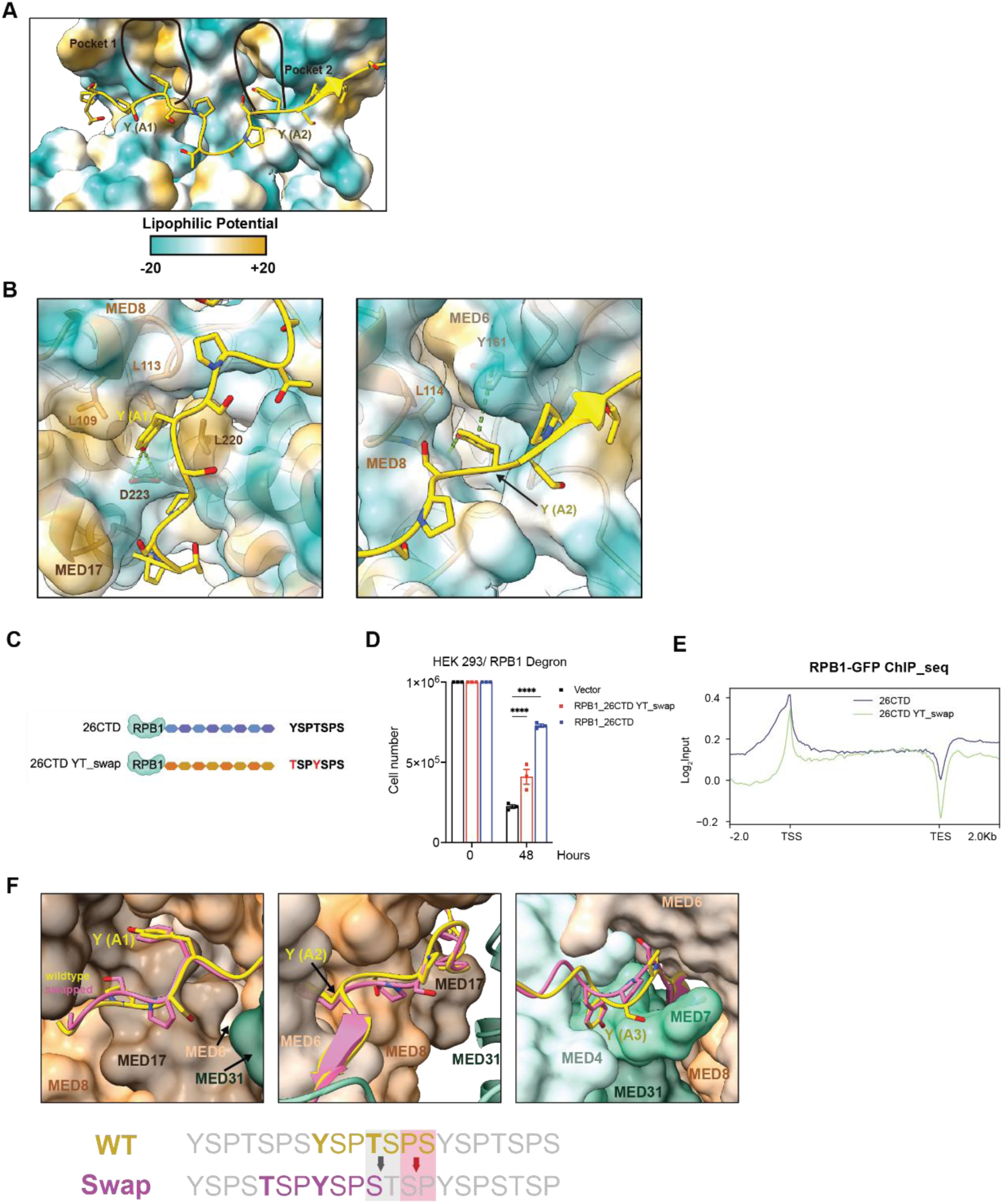
Functional flexibility in Ser2 and Ser5 positioning within CTD heptad. (A) Structure of the yeast Mediator (PDB Code: 8CEO) bound to the RNAPII CTD. The Mediator is shown as a surface representation and is colored by lipophilic potential, while the Pol II CTD is shown as a cartoon. (B) Interaction between region A of the CTD with the yeast Mediator head subunits (PDB Code: 8CEO). The Mediator subunits are shown as cartoons with a transparent surface representation, which is colored by electrostatic surface potential. Relevant Mediator side chains that comprise each hydrophobic pocket or interact with the hydroxyl group of the CTD tyrosine are shown as sticks. Region A of the CTD is shown in yellow. (C) Schematic of RPB1 26CTD and 26CTD YT_swap. (D) Cell density after transfection of RPB1 26CTD or 26CTD YT_swap constructs in HEK 293 RPB1 degron cells after auxin administration. n = 3 independent experiments (mean ± SEM); ****p < 0.0001, comparison was performed using two-way ANOVA with Tukey correction. (E) Merged metagene plot of RPB1 ChIP-seq data with 26CTD and 26CTD YT_swap constructs. (F) Region A of the wild-type CTD (yellow) or a model of the swapped CTD (pink) bound to the yeast Mediator. Mediator head subunits are colored varying shades of orange, while Mediator middle subunits are colored varying shades of green.

Together, these structural features highlight why Tyr1 is indispensable for Mediator recognition: the bulky aromatic ring anchors the CTD into regularly spaced hydrophobic pockets, while the hydroxyl group fine-tunes binding through hydrogen bonds. The periodicity of these pockets—matching the seven–amino acid spacing of the CTD repeat—further explains why altering CTD length is detrimental. Modeling hexameric or octameric repeats onto the yeast Mediator produced severe steric clashes (Figure SI 4B, 4C), consistent with the lethal phenotypes of repeat length changes observed in yeast (Nonet and Young 1989; West and Corden 1995).

### Flexibility in Ser2 and Ser5 positioning within the CTD heptad

Another strictly conserved feature of the CTD heptad is the pair of SP motifs, since phosphorylation of Ser2 and Ser5 is central to transcription regulation (Ho and Shuman 1999). Ser5 phosphorylation promotes promoter escape and recruits mRNA capping enzymes (Lu et al. 1992; McCracken et al. 1997; Cho et al. 1997), while Ser2 phosphorylation accumulates across the gene body and peaks near termination sites to recruit elongation and termination factors (Barillà et al. 2001; Kim et al. 2004; Meinhart and Cramer 2004). Given these distinct roles, we asked how crucial the precise positioning of Ser2 and Ser5 within the heptad is for CTD function.

In the canonical sequence (Y_1_S_2_P_3_T_4_S_5_P_6_S_7_), Ser2 is flanked by Tyr1 and Ser5 by Thr4. To test whether these contexts matter, we swapped the positions of Tyr1 and Thr4, generating a “swapped” motif (**<u>T_1_</u>**S_2_P_3_**<u>Y_4_</u>**S_5_P_6_S_7_) while preserving overall amino acid composition and heptad spacing (Figure 4C). Using RPB1 constructs with 26 repeats which support viability, albeit with slightly slower growth than a full-length 52CTD (Figure SI 4D) (Sawicka et al. 2021; Q. Zhang et al. 2024), we found that the Ser2/Ser5 swap also sustained viability, though with moderately reduced growth in both the degron and α-amanitin systems (Figure 4D, SI 4D).

ChIP-seq analyses on cells expressing the swapped CTD revealed that mutant Pol II localized to promoters, although with a reduced enrichment at transcription start sites (TSS) (Figure 4E). Structural modeling provides a rationale for this tolerance. In both the wild-type and swapped CTDs, Tyr1 residues, which are key anchors for Mediator binding, remain in register within the seven-residue repeat. The principal difference is that the Pro6–Ser7 motif of the wild-type sequence becomes a Ser–Pro motif preceding Tyr4 in the swapped version (Figure 4F). Energy minimization of the swapped CTD within the Mediator interface revealed only minor deviations from the wild-type fit (Figure 4F, SI 4E). For instance, in region A, the flipped Ser–Pro motif introduces a polar serine into a non-lipophilic pocket and shifts the adjacent proline, but the alteration is structurally tolerated (Figure SI 4F). Thus, while Ser2 and Ser5 phosphorylation sites are indispensable, their exact placement within the heptad is flexible. The CTD can accommodate positional swaps without abolishing Mediator binding or transcription, explaining the retention of viability with only modest transcriptional defects.

### Ser2/Ser5 Positional Swap Does Not Impair CTD Phosphorylation or promoter clearance

Having shown that the Ser2/Ser5 positional swap still allows PIC assembly, we next asked whether it affects CTD phosphorylation by transcriptional kinases such as TFIIH and P-TEFb. We purified a CTD construct containing 26 swapped repeats and assessed its phosphorylation by an electrophoretic mobility shift assay (EMSA). Upon kinase incubation, the swapped CTD showed progressive gel shifts consistent with phosphorylation (Figure 5A). Although migration was slightly slower for the swapped CTD compared to the wild-type 26CTD, both were efficiently phosphorylated. Kinetic analysis of TFIIH activity confirmed comparable catalytic efficiency for swapped and wild-type CTDs (Figure 5B).

**Figure 5:**
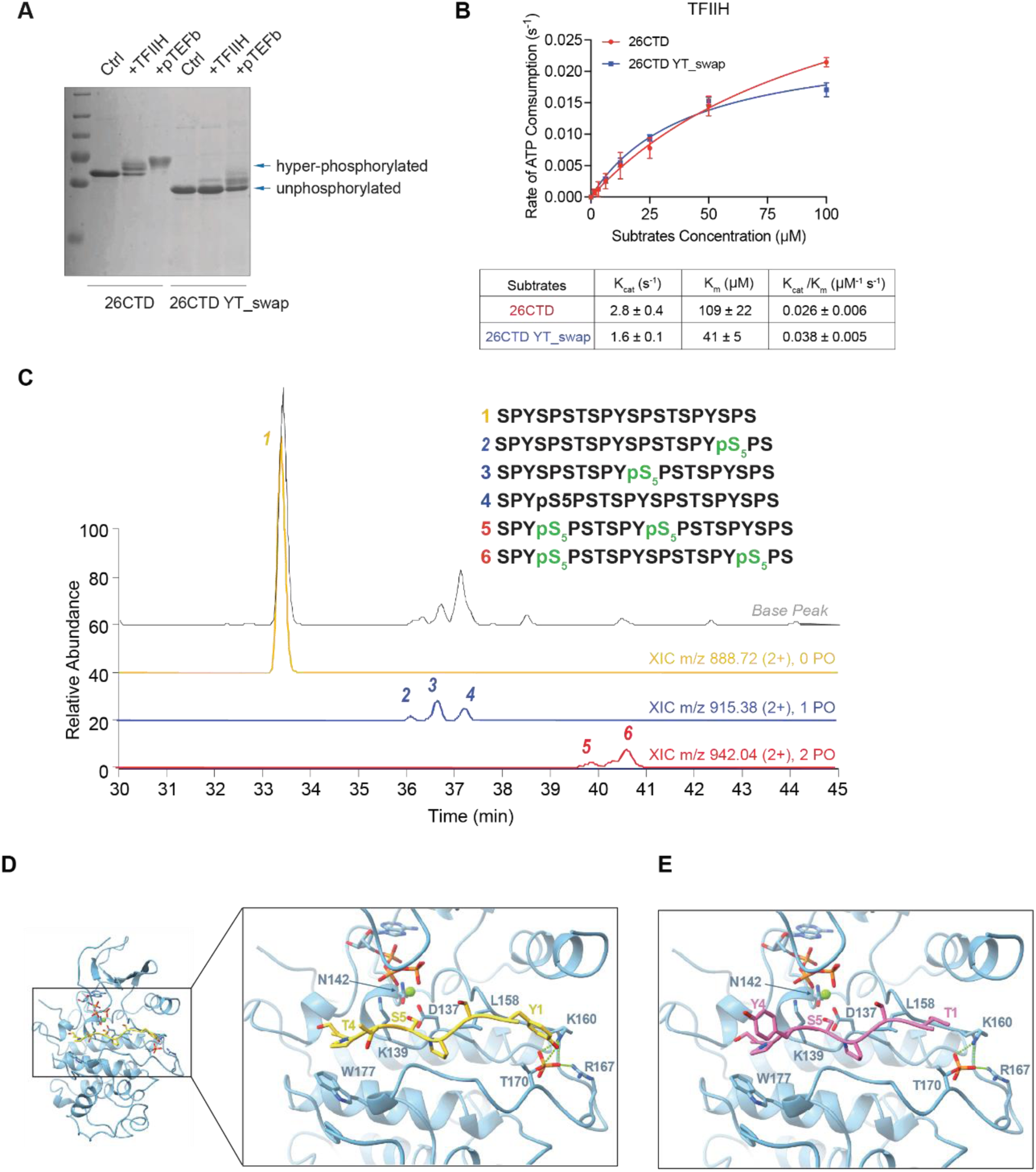
Ser2/Ser5 Positional Swap Does Not Impair CTD Phosphorylation or PIC Disassembly. (A) SDS-PAGE gel analysis of unphosphorylated CTD samples treated with TFIIH and pTEFb. Change in electrophoretic mobility caused by phosphorylation is shown by gradual shift of protein bands corresponding to GST-26CTD and GST-26CTD YT_swap. (B) Kinase activity assay of wild-type 26CTD (red) and 26CTD YT_swap (dark blue) by TFIIH fitted to the Michaelis-Menten kinetic equation. The Michaelis-Menten kinetic parameters kcat/Km (mM-1 s-1) are given for each respective fit. Each measurement was conducted in triplicate, with standard deviations shown as error bars. (C) Extracted ion chromatographic traces (XIC) for the LC-MS/MS analysis for kinase specificity toward the YT_swap CTD sequence. The gold LC trace corresponds to the unphosphorylated peptide, whereas the blue trace indicated the monophosphorylated species with peak numbers matching the sites of phosphorylation indicated on the sequences above the LC traces. The red-colored LC traces correspond to the di-phosphorylated species. (D) Model of CDK7 bound to the Pol II CTD (yellow). Ser5 was positioned for phosphorylation within the CDK7 active site. (E) Model of CDK7 bound to the swapped Pol II CTD (pink). Ser5 was positioned for phosphorylation within the CDK7 active site.

To determine whether the residue swap alters the specific site of phosphorylation, we conducted high-resolution mass spectrometry to map phosphorylation sites within the swapped CTD (Figure 5C and SI 5A). Given the analytical challenge posed by the repetitive CTD sequence, we engineered a simplified construct containing three heptad repeats fused to GST (Figure 5C). After kinase treatment with TFIIH, the CTD fragment was released via 3C protease digestion and analyzed by liquid chromatography (LC) coupled with ultraviolet photodissociation (UVPD) mass spectrometry for single-residue resolution of phosphorylation sites. The LC-MS trace of singly phosphorylated species revealed three well-resolved peaks, each corresponding to a different heptad. UVPD analysis confirmed that in all cases, phosphorylation occurred at Ser5 (Figure 5C and SI 5A), demonstrating that TFIIH continues to preferentially target Ser5 even when its flanking residues are altered. Additional peaks corresponding to doubly phosphorylated species also showed consistent Ser5 phosphorylation across multiple repeats (Figure 5C). Altogether, these data confirm that Ser5 remains the dominant phosphorylation site in the swapped CTD, despite the altered sequence context. This selectivity is consistent with CDK7’s structural determinants of Ser5 recognition (Ramani et al. 2020): a conserved Trp stacks with the CTD proline, and Tyr1 favors interactions with the phosphorylated T-loop. These interactions are minutely affected in the swapped CTD but are disrupted when Ser2 occupies the catalytic position, explaining TFIIH’s persistent Ser5 preference (Figures 5D and 5E, SI 5B).

Altogether, these data demonstrate that the swapped CTD preserves essential features for recognition and phosphorylation by TFIIH, enabling Pol II to undergo hyperphosphorylation and transition into elongation.

### Ser2/Ser5 swapping affects transcription elongation due to defective regulator recruitment

Heatmap analysis of ChIP-seq data revealed that the Ser2/Ser5 swapped mutant showed an obvious increase in Pol II occupancy across gene bodies, indicating a difference in Pol II distribution (Figures 6A, 6B). To assess how these changes affect transcriptional output, we performed spike-in–normalized TT-seq in HEK293 RPB1 degron cells transiently expressing either swapped or wild-type 26CTD. Endogenous RPB1 was degraded using the OsTIR1–auxin system, and nascent RNA was labeled by 4-thiouridine (4sU) incorporation. Heatmaps and metagene profiles revealed a broad reduction in sense and antisense transcription in the swapped CTD group (Figures 6C, SI 6A-C). An MA plot confirmed a global downregulation of nascent RNA, with 37.7% of transcripts significantly reduced in the swapped condition (adjusted p < 0.05) (Figure 6D). These results show that Ser2/Ser5 swapping leads to widespread impairment of nascent RNA synthesis.

**Figure 6:**
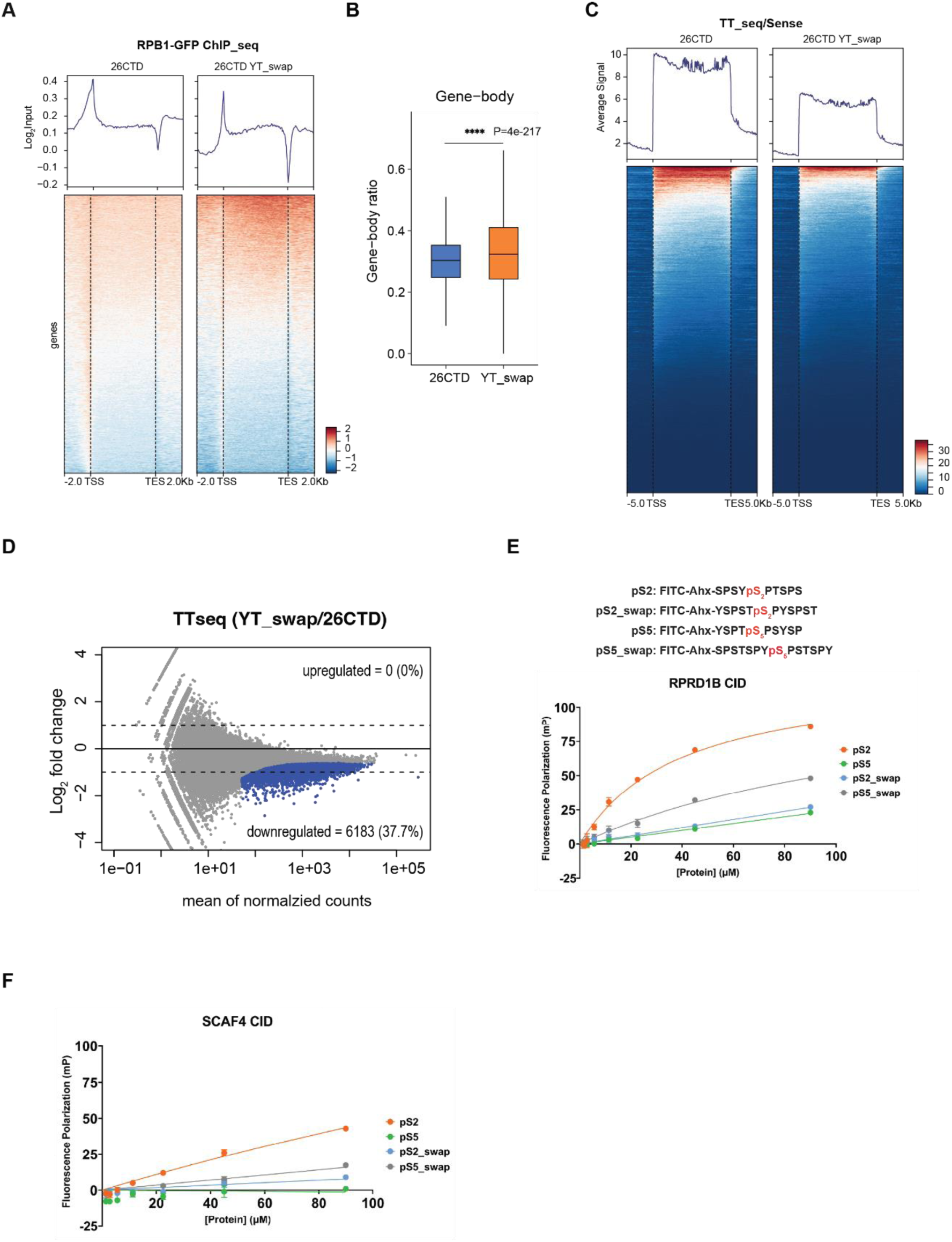
Ser2/Ser5 swapping affects transcription elongation and 3’ end processing. (A) Input normalized metagene plots (top) and heatmaps (bottom) from ChIP-seq analysis of RPB1 26CTD and RPB1 26CTD YT_swap. Two biological replicates (n = 2). (B) Box plot showing the distribution of gene-body signal ratio (GB / (PR + GB + DS)) for yCTD and swap Pol II ChIP-seq samples. Statistical significance was calculated using a paired Wilcoxon signed-rank test. PR: promoter region (TSS to ±300 bp), GB: gene bodies (TSS+500 bp to TES), DS: downstream region (TES+2 kb). (C) Average sense signal and heatmaps in 26CTD and YT_swap CTD for TT-seq, 2 replicates for each group. (D) MA plot analysis of the differentially expressed genes in YT_swap CTD compared with 26CTD. (E-F) Fluorescence polarization (FP) measurement of the binding of pSer2 or pSer5 FITC-labeled CTD or YT_swapped CTD peptides containing two repeats to RPRD1B (E) and SCAF4 (F). Data were fitted to a 1:1 binding model. Binding assays were performed in triplicate. Error bars indicate the standard deviation of the mean.

We next asked why the swapped mutant seemed to stall Pol II in gene body and attenuate elongation. Although many CTD readers can recognize both pSer2 and pSer5, some show strong positional preference. CID-containing proteins, which function in elongation, 3′ end processing, and termination, generally prefer pSer2 (Pineda et al. 2015; Jasnovidova et al. 2017; Moreno et al. 2025). For example, RPRD1B, a 3′ end processing factor, binds tightly to a pSer2 CTD but not to a pSer5 CTD (Moreno et al. 2025; Ni et al. 2014), while SCAF4 prefers a doubly phosphorylated CTD (pSer2/pSer5) but shows only weak binding to either site individually (Gregersen et al. 2019). To test how Ser2/Ser5 swapping affects these interactions, we measured binding of FITC-labeled CTD peptides (two heptads in length) to RPRD1B and SCAF4 (Figures 6E, 6F). RPRD1B binding was strongly reduced (>3-fold) for the swapped pSer5 CTD compared to pSer2, with no detectable binding to wild-type pSer5 or swapped pSer2. The swapped CTD caused almost no detectable binding to SCAF4.

Thus, Ser2/Ser5 swapping compromises recognition of SP motifs by key CTD-binding proteins, providing a mechanistic explanation for the elongation and 3′ end processing defects of the swapped CTD Pol II.

## DISCUSSION

In our study, we utilized multi-disciplinary methods with newly designed artificial CTD constructs to explore why evolution strongly favors the repetitive heptad motif in the CTD of Pol II. The conserved YSPTSPS sequence has long been a focal point of speculation. In 2008, Chapman et al. proposed that the CTD heptad evolved from shorter YSPX or SPXY motifs, converging on a consensus of SPXYSPX centered around tyrosine (Chapman et al. 2008). Our structural and bioengineering studies advanced this effort by dissecting the contribution of CTD length, understanding the CTD spacing and recognition motifs, and providing a molecular rationale for a conserved “CTD grammar”.

### Roles of the CTD at Different Stages of Transcription

The striking periodicity and evolutionary conservation of the Pol II CTD has long fascinated transcription researchers. Our data, along with previous genetic studies, reveal mechanistic insights into how the CTD contributes to distinct stages of Pol II-mediated transcription. During PIC formation, the unphosphorylated CTD wraps around the interface between the head and middle modules of the Mediator complex (Figure 7A). We discovered that tyrosine residues function as anchors by inserting into evenly spaced hydrophobic pockets along the Mediator interface between the head and middle modules, analogous to nails driven into wood (Figures 4A). This extensive stretch of heptad repeats must remain unphosphorylated to maintain a stable PIC. Our structural predictions and functional assays using RPB1 variants with altered CTD lengths indicate that a minimum of 17 repeats is required to span the Mediator interface to fulfill the necessary architectural role and to ready Pol II for subsequent phosphorylation (Figures 1D). Consistent with this, super-resolution imaging confirms that CTD length directly influences PIC assembly in vivo.

**Figure 7:**
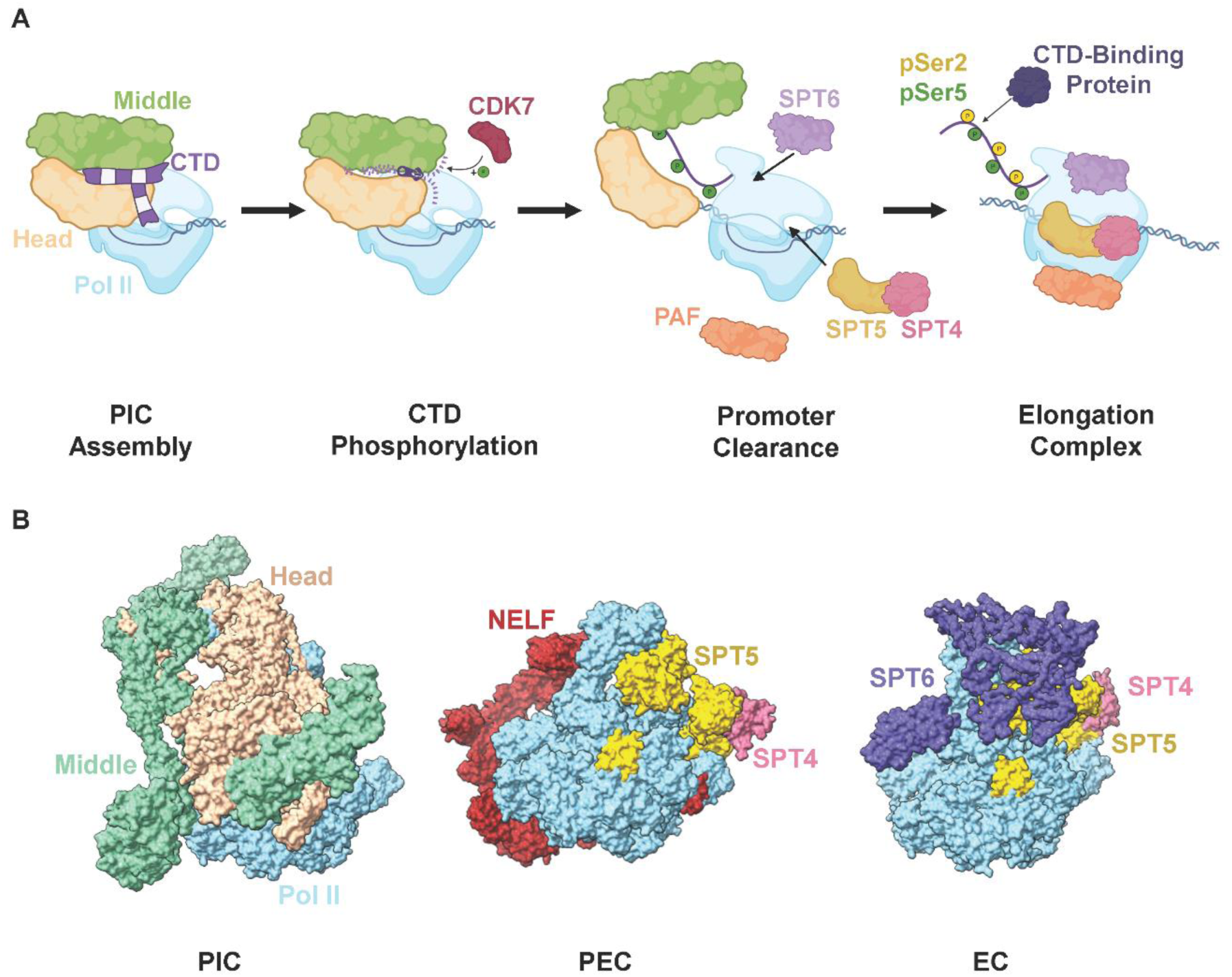
Roles of the CTD at Different Stages of Transcription. (A) Zipper model of RNAPII in transcription. (B) Surface representation of Pol II in complex with the Mediator Head and Middle (left, PDB Code: 8XGS), NELF and DSIF (middle, PDB Code: 8UHD), or SPT6 and DSIF (right, PDB Code: 6GMH). Pol II is shown in the same orientation for each structure in the figure.

Following transcription initiation, phosphorylation of the CTD is essential for promoter clearance. Ser2 and Ser5 residues become phosphorylation targets for SP-specific kinases, which promote promoter clearance by progressively adding negative charges to the CTD heptads. It is the cumulative charge density, rather than the exact phosphorylation position, that is critical for disrupting CTD-Mediator interactions. This modification proceeds in a “zipper-like” manner: heptad repeats are sequentially phosphorylated, advancing towards Mediator interface. As negative charges build, they weaken the hydrophobic anchoring of the CTD, ultimately dislodging Pol II from the Mediator and triggering promoter clearance (Figure 7A). Mediator release is a prerequisite for Pol II to engage DSIF, which binds to the same Pol II surface as the Mediator near the stalk region (Figures 7B) (Su and Vos 2024). Once freed from Mediator, Pol II can form a stable pre-elongation complex (PEC) and subsequently assemble elongation factors such as SPT6 and PAF1 (Figures 7B) (Vos et al. 2018). This zipper model highlights that PIC disassembly is not a sudden, stochastic event but a coordinated, directional process driven by kinase activity. Thus, the CTD has dual roles in early transcription—stabilizing the PIC in its unphosphorylated state and promoting promoter clearance through phosphorylation to prime Pol II for productive elongation.

While the general requirement for phosphorylation emerges during initiation, the precise positioning of the phospho-sites only becomes critical during elongation. Specifically, swapping the positions of Ser2Pro3 and Ser5Pro6 motifs reduces elongation efficiency because phospho-Ser2 serves as a key recruitment site for transcription regulatory factors, and scrambling the heptad sequence disrupts this functional specificity. Consequently, mis-patterned CTDs impair the recruitment of elongation factors and other 3’ end regulatory proteins, leading to defects in later stages of transcription.

Overall, a CTD sequence motif of the form a XSPXXSP, where one “X” is tyrosine and the others are small uncharged residues, can support Pol II function and cell viability. For effective transcription initiation, a minimum of 1/3 of continuous endogenous CTD repeats must conform to this “grammar”, although the precise position within the flexible CTD scaffold appears less critical.

Our findings are consistent with the emerging role of the CTD in liquid-liquid phase separation (LLPS) and transcriptional condensate formation (Rippe and Papantonis 2025). Recent studies have demonstrated that unphosphorylated Tyr1 residues drive LLPS of the CTD with Mediator and other transcription factors, concentrating Pol II and regulatory components into dynamic condensates that enhance efficiency of PIC assembly (Zhang et al. 2024; Lyons et al. 2025). In this context, the zipper model posits that the periodic Tyr1-Mediator hydrophobic interactions not only stabilize the PIC scaffold but also serve as multivalent anchors that nucleate and propagate LLPS, with a minimal stretch acting as a critical “seed” length to achieve sufficient valency for phase separation (Ling et al. 2024). Progressive phosphorylation of adjacent SP motifs, by introducing electrostatic repulsion and steric hindrance, progressively disrupts these hydrophobic contacts, effectively “unzipping” the CTD from the Mediator interface and ejecting Pol II from the PIC condensate, enabling promoter escape and Pol II handover to elongation factors like DSIF and NELF. This phosphorylation-triggered transition from a condensed, initiation-competent state to a more diffusive, elongating conformation reconciles the CTD’s dual roles as both a condensate organizer and a dynamic release switch, highlighting how its low-complexity grammar encodes spatiotemporal control over transcription through coupled structural and phase behaviors.

### Interpreting Genetic Outcomes of CTD Variants

Our proposed CTD grammar provides a framework to interpret previous mutational studies of the Pol II CTD. First, it explains why a minimum number of heptad repeats is required for cell viability, even though the CTD doesn’t affect the polymerase catalytic activity (Zehring et al. 1988). The CTD must span the Mediator interface and establish tyrosine-mediated hydrophobic contacts that stabilize the PIC. Mutations that retain a sufficient stretch of the consensus YSPTSPS motifs generally support cell survival (West and Corden 1995; Sawicka et al. 2021; Meininghaus and Eick 1999; Zhang et al. 2024; Nonet and Young 1989). In humans, approximately 17 repeats (about 1/3 of full-length) are sufficient for Mediator binding and CTD phosphorylation. However, artificially engineered CTDs with alternating blocks—such as one-quarter of the repeats containing the native tyrosine, followed by one-quarter with phenylalanine replacing tyrosine, and so on—fail to support human cell growth, despite an overall motif preservation of 50% (Shah et al. 2018). This underscores the importance of not just the presence but also the continuity of functional motifs.

Second, the periodical spacing of the hydrophobic pockets along the Mediator head-middle interface explains why CTD variants with altered repeat length (e.g., octad repeats) are nonviable in *S. cerevisiae* (Stiller and Cook 2004). Misalignment disrupts Tyr1 anchoring, leading to impaired PIC assembly. This also explains why tyrosine is the least varied residue in the CTD; even a conservative mutation to phenylalanine in every heptad causes lethality in *S. cerevisiae* (West and Corden 1995) and in humans (Descostes et al. 2014), highlighting the need for an interaction with both the hydrophobic ring and the hydroxyl group of Tyr1.

Third, the requirement for uncharged residues reflects the hydrophobic environment during PIC formation. Even seemingly minor substitutions (such as replacing the seventh residue with charged residues like Glu or Lys) disrupt these interactions and displace Pol II localization as detected by ChIP-seq (Schwer and Shuman 2011; West and Corden 1995; Kecman et al. 2018; Zhang et al. 2012; 2024). Similarly, RPB1 variants with a minimal length of consensus repeats can support cell growth, whereas RPB1 variants that lack enough uncharged sequential repeats cannot support cell growth due to the inclusion of charged residues (West and Corden 1995). Our engineered hybrid CTDs demonstrate that transcription proceeds as long as a continuous intact stretch conforming to the CTD grammar is preserved. While the exact positioning of the uncharged repeats within the CTD is less critical, there is a slight preference for cell fitness when the conserved heptads are in the proximal region of the CTD.

Finally, the SP motif is indispensable for Pol II to exit the PIC. Our “zipper model” proposes that progressive phosphorylation of SP motifs sequentially disengages CTD contacts with Mediator, prompting promoter clearance. This is consistent with the substrate specificity of transcriptional kinases (TFIIH/CDK7, P-TEFb/CDK9, CDK12, CDK13), all of which target SP motifs, especially Ser5-Pro6 (Mayfield et al. 2019; Czudnochowski et al. 2012). The absolute conservation of prolines further underscores their essential structural role, as mutation of either proline is lethal (Schwer and Shuman 2011).

## Limitation

While our model accounts for the majority of known CTD mutations, certain species-specific exceptions remain. For instance, a Tyr1-to-Phe substitution is tolerated in *S. pombe*, but not in most other systems (Schwer and Shuman 2011; West and Corden 1995; Descostes et al. 2014; Shah et al. 2018). We propose that this reflects subtle species-specific Mediator–CTD interactions. In *S. pombe*, the hydroxyl group of tyrosine may be dispensable for Mediator binding, whereas it is essential in other organisms. This notion is supported by co-evolutionary evidence of the CTD and Mediator complex: mutations in Mediator subunits can sometimes suppress CTD defects (Yuryev and Corden 1996). Another notable unexplained mutation is Thr4-to-Ala, which is lethal in humans but tolerated is in yeast (Schwer and Shuman 2011; Hintermair et al. 2012). Our data suggest that defective Thr4 phosphorylation impairs transcription termination of critical cell cycle regulators, leading to cell death in human cells (Moreno et al. 2024).

## Conclusion: The Zipper Model of CTD Function

Taken together, our genetic, biochemical, and cellular evidence converge on a “zipper model” for CTD function (Figure 7A). In its unphosphorylated state, the CTD is largely uncharged and threads through the Mediator complex, with tyrosines anchoring into hydrophobic pockets to stabilize the PIC. Upon transitioning into transcription initiation, phosphorylation of SP motifs introduces negative charges that progressively “unzip” the CTD from the Mediator, triggering promoter clearance. As elongation proceeds, Ser2 phosphorylation accumulates, recruiting 3’ end processing factors and other elongation/termination regulators. This dual role of the CTD—as a scaffold for PIC assembly and a dynamic platform for regulatory factor recruitment—has driven its co-evolution with the Mediator.

## METHODS

### Cell lines, Reagents, and Antibodies

HEK293T and U2OS cells were obtained from ATCC. The inducible HEK293 RPB1 degron cell line was generously provided by Dr. Robert G. Roeder. None of the cell lines used in this study are listed in the databases of commonly misidentified cell lines maintained by the International Cell Line Authentication Committee (ICLAC) and NCBI BioSample. All cell lines were cultured in DMEM supplemented with 10% fetal bovine serum (FBS) and 1% penicillin-streptomycin and incubated at 37°C with 5% CO2.

Transfections were carried out using Polyethyleneimine (PEI; Polysciences) or FuGENE HD (Promega) according to the manufacturers’ protocols. For endogenous RPB1 degradation, α-amanitin (Sigma) was used at a final concentration of 2.5 μg/mL for a total of 72 hours. In HEK293 RPB1 degron cells, cells were treated with doxycycline (2 μg/mL) to induce OsTIR1 expression for 12 hours, followed by treatment with auxin (500 μM) for 48 hours to trigger targeted degradation of RPB1-AID.

Anti-GFP antibody (Proteintech, catalog #50430-2-AP; 5 μg/sample) was used for ChIP experiments. Anti-MED1 antibody (Abcam, ab313323, 1:100 dilution) was used for immunofluorescence staining. The anti-rabbit IgG antibody was purchased from Invitrogen (catalog #08-6199) as a control.

### CTD engineering

All 26x yCTD constructs (WT, S5E, T4E or S7E in every other repeat, T4E in every repeat, and YT swapped CTD) were synthesized by GenScript. Constructs containing different numbers of CTD repeats (3CTD, 9CTD, 13CTD, 15CTD, 17CTD, 20CTD) were generously provided by Dr. Carl Wu. For mammalian protein expression, YFP-RPB1aAmr-WT (52CTD) was obtained from Addgene. YFP-RPB1aAmr-different CTD repeats, YT swapped CTD, S5E/T4E/S7E mutants, and hybrid mutants were generated via PCR-based cloning using the ClonExpress II One Step Cloning Kit (Vazyme). All coding sequences were confirmed by DNA sequencing.

### Structural Modeling

Because the yeast PIC structure (PDB Code: 8CEO) was captured at a lower resolution than the human PIC structures (PDB Codes: 8GXS, 8GXQ), we generated a model of the human PIC (PDB Code: 8GXS) bound to the yeast CTD repeats. To position the yeast repeats into the human PIC we aligned the yeast PIC structure to the human PIC structure using MED6 as the reference chain for both structures. Then we energy minimized the yeast CTD repeats within the human PIC by applying the OLS-2005 force filed in the MAESTRO (version 14.2) from the Schrodinger suite (Shivakumar et al. 2010; Kaminski et al. 2001; Schrödinger, LLC. 2025a). The final model and all structure figures were visualized in ChimeraX (version 1.9) (Goddard et al. 2018).

To generate a model of the yeast PIC bound to a phospho-mimetic CTD, we mutated the wild-type CTD to the mutant T4E CTD in MEASTRO (PDB Code: 8CEO), and then we energy minimized the mutant CTD repeats within the yeast PIC by applying the OLS-2005 force filed in MAESTRO (Shivakumar et al. 2010; Kaminski et al. 2001; Schrödinger, LLC. 2025a).

We used PDB ePISA (European Bioinformatics Institute 2007; Krissinel and Henrick 2007) to calculate the interaction area between Mediator subunits with and without the CTD present. PyMOL (version 3.1.3) (Schrödinger, LLC. 2025b) was used to measure the length of CTD repeats in the yeast and human PIC complex structures to generate an average CTD heptad length.

To compare Pol II’s interaction with the Mediator to the interaction between Pol II and pausing or elongation factors, we aligned the human PIC structure (PDB Code: 8XGS) to a structure of Pol II in complex with NELF and DSIF (PDB Code: 8UHD), or a structure of Pol II in complex with SPT6 and DSIF (PDB Code: 6GMH), using RPB1 as the reference chain.

### Cell growth assay

For the α-amanitin system, HEK293T cells were seeded into 6-well plates at a density of 5 × 10⁵ cells/mL and transfected with RPB1aAmr-YFP containing various CTD constructs. After 24 hours of transfection, α-amanitin (Sigma) was added at a final concentration of 2.5 μg/mL, with fresh medium containing α-amanitin replenished daily. After 72 hours, cell numbers were counted using a cell counter with trypan blue staining.

For the RPB1 degron system, inducible HEK293 RPB1 degron cells were seeded into 6-well plates at a density of 5 × 10⁵ cells/mL and transfected with RPB1aAmr-YFP containing various CTD constructs. After 36 hours of transfection, doxycycline (2 μg/mL) was added to induce OsTIR1 expression. 12 hours later, auxin (500 μM) was added to induce RPB1-AID degradation. Cell numbers were determined after an additional 48 hours using a cell counter with trypan blue staining.

### STORM imaging

Dual-color STORM imaging was performed using a Nikon Eclipse-Ti2 inverted microscope equipped with a 100× (N.A. 1.45) Plan Apo oil-immersion objective (Olympus) and an EM-CCD camera (Andor iXon Life 897). Four laser lines, including 405 nm (Coherent OBIS 140 mW), 488 nm (Coherent Sapphire 300 mW), 560 nm (MPB Communications, 1500 mW), and 642 nm (MPB Communications, 2000 mW), were coupled into the microscope via the rear port. For illumination, the laser beams were laterally offset at the objective’s back aperture to produce near-total internal reflection (TIR), selectively exciting fluorophores within a narrow axial region close to the coverslip (Zhou et al. 2019).

For dual-color experiments, AF647 and CF583R fluorophores were imaged sequentially. Imaging was conducted in an enzymatic oxygen-scavenging buffer composed of 100 mM Tris-HCl (pH 7.5), 100 mM cysteamine, 5% (w/v) glucose, 0.8 mg/mL glucose oxidase, and 40 µg/mL catalase (all from Sigma-Aldrich). During acquisition, the 560-nm or 642-nm lasers were applied continuously at ∼2 kW/cm² to drive most fluorophores into long-lived dark states, while the 405-nm laser was used at low adjustable intensity (0-1 W/cm²) to stochastically reactivate single emitters. Approximately 60,000 frames (split evenly between the two color-channels) were recorded per dataset at 110 Hz frame rate.

Post-acquisition processing involved localizing individual fluorophores and rendering super-resolution images by plotting each localization as a 2D Gaussian. Channels alignment was achieved using fiducial markers (Thermo Fisher Scientific fluorescent microspheres, F8803) that fluoresced in both channels to ensure accurate channel overlay.

### STORM data analysis

Image analysis and quantification were performed using custom MATLAB scripts applied to dual-color STORM datasets. Regions of interest (ROIs; ∼2 µm × 2 µm) containing MED1 puncta were selected from reconstructed images. Within each ROI, the largest MED1 and RPB1 clusters were segmented using adaptive thresholding and binary masking. A composite mask was generated to capture either colocalized regions or, in the absence of overlap, the MED1 cluster alone.

To compare the MED1 cluster sizes across experimental conditions, the area of each MED1 cluster was measured within its respective mask and then normalized to the mean MED1 cluster area measured from the 52CTD control. To determine the percentage of MED1 clusters colocalized with RPB1 clusters, segmented MED1 clusters were classified as either “colocalized” if they overlapped with an RPB1 cluster, with overlap defined as any nonzero intersecting area between the two binary masks. Colocalization percentage was calculated by dividing the number of colocalized MED1 clusters by the total number of MED1 clusters analyzed.

To assess spatial correlation between MED1 and RPB1 signals at the pixel level, Pearson’s correlation coefficients (PCC) were calculated for each ROI. Pixel intensities were extracted from the MED1 (CF583R) and RPB1 (AF647) channels and compared using the standard PCC formula. In addition, to quantify the degree of spatial overlap, the percentage of each MED1 cluster area overlapping with the corresponding RPB1 signal was calculated by dividing the overlapping area by the total MED1 cluster area.

All measurements were aggregated across multiple ROIs for each construct. Statistical analyses were performed to evaluate differences among constructs.

### Electrophoretic Mobility Assay

TFIIH or pTEFb kinase reactions were prepared by incubating 1 mg/mL GST-26CTD or GST-26CTD YT_swap with 0.1 mg/mL pTEFb or 0.0025 mg/mL TFIIH in a reaction buffer containing 50 mM Tris-HCl at pH 8.0, 50 mM MgCl_2_ and 2 mM ATP. Kinase reactions and no kinase control reactions were incubated at 30°C for 16 h. Reactions were quenched by addition of 2X Laemmli SDS buffer in preparation for SDS-PAGE analysis. Kinase reaction samples were boiled at 95°C for 5 mins and then run on a 15% denaturing Tris-glycine polyacrylamide gel, which was stained with Coomassie solution after running.

### Kinase activity assay

The TFIIH kinetic activity assay was performed in a 10 μL reaction volume containing 0-100 μM of the substrate (GST-26CTD or GST-26CTD YT_swap) in a reaction buffer containing 50 mM Tris at pH 8.0, 20 mM MgCl_2_, and 50 μM ATP. The reaction was initiated by adding 20 nM of kinase and incubating at 30°C for 30 minutes before quenching with 10 μL of room temperature Kinase-Glo Detection Reagent (Promega). The mixtures were incubated at room temperature for 10 minutes with the reagent before obtaining luminescence readings in a Tecan plate reader 200. The readings were translated to ATP concentration using an ATP standard curve determined with the Kinase-Glo Detection Reagent. Kinetic data were obtained in triplicate and fitted to the Michaelis-Menten kinetic equation to obtain respective kinetic parameters *k_cat_* (min-1) and *K_m_*(μM) in GraphPad Prism 9.

### UVPD Tandem Mass Spectrometry and Data Analysis

The 3X swapped CTD construct was analyzed using identical LC–UVPD-MS workflows as described previously (Mayfield et al. 2019; 2017; Irani et al. 2019; Burkholder et al. 2019; Ramani et al. 2020).

0.2 *μ*M TFIIH was used to treat 1 mg/mL of a GST-tagged swapped CTD substrate that contains only 3 CTD repeats for 12 hours in a reaction buffer containing 2 mM ATP, 50 mM Tris pH 8, and 10 mM MgCl_2_. The reaction was quenched by addition of 10 mM EDTA. The GST tag was cleaved from the 3X CTD peptide by digesting the sample with 3C-protease at a molar ratio of 100:1 protein/protease. The digested sample was desalted in a C18 spin column and resuspended in 2% acetonitrile and 0.1% formic acid in HPLC-grade water for LC–MS analysis.

A Dionex Ultimate 3000 nano liquid chromatography system (Thermo Scientific) plumbed for direct injection into a 75 *μ*m ID Picofrit analytical column (New Objective, Woburn, MA) was used to separate the peptide sample. 1 μL was injected for separation using a 1.8 *μ*m UChrom C18 analytical column (NanoLCMS Solutions, Oroville, CA) packed in-house to 20 cm. Mobile phase A was composed of HPLC-grade water containing 0.1% formic acid and mobile phase B was composed of HPLC-grade acetonitrile containing 0.1% formic acid. Separation was carried out using a gradient that was optimized for the specific sample, at a flow rate of 0.300 *μ*L/min. An Orbitrap Fusion Lumos Tribrid mass spectrometer (Thermo Fisher Scientific, San Jose, CA) using a NanoFlex electrospray source was used to analyze eluted peptides in positive polarity mode. As described previously, the mass spectrometer was fitted with an excimer laser operated at 193 nm (Coherent, Santa Clara, CA) and was modified to allow for UVPD in the dual linear ion trap (Klein et al. 2016). Spectra were acquired in the Orbitrap mass analyzer using resolution settings of 60 K for MS1 events and 30 K (at *m*/*z* 200) for MS/MS events. Two laser pulses of 1.5 mJ for UVPD in the low-pressure ion trap were used to activate target peptides (Klein et al. 2016).

MS/MS spectra were deconvoluted using the Xtract algorithm within XCalibur QualBrowser, applying a signal-to-noise threshold of 3. Fragment ions were assigned using ProSight Lite by matching to nine characteristic ion types generated by UVPD of peptides (a, a+1, b, c, x, x+1, y, y–1, z). Phosphosite localization was achieved by computationally adding the mass of a phosphate group (+79.97 Da) to each candidate Thr or Ser residue and identifying fragment ions that retained the modification. Relative abundances of phosphorylated species were quantified from the LC peak area corresponding to each eluting peptide. The ion current of each phosphopeptide precursor was integrated across its elution profile and normalized to the total ion current of all phosphopeptide isomers. Because isobaric peptides share identical chemical composition, ionization efficiencies were assumed to be equivalent across species. Abundance comparisons were restricted to phospho-isomers of the same peptide sequence. All experiments were performed with at least two biological replicates, and mass spectra were validated both manually and via commercial search algorithms.

### CID Domain Purification

BL21 (DE3) cells expressing either 6XHis-GST-RPRD1B-CID were grown in 4 liters of Luria-Bertani (LB) broth containing 50 μg/mL kanamycin at 37 °C or 6XHis-HGST-SCAF4-CID were grown in 2 L of Terrific broth (TB) containing 50 μg/mL kanamycin at 37 °C. Once the LB cultures reached an OD_600_ of ∼0.6, protein expression was induced by adding 0.5 mM isopropyl-β-d-thiogalactopyranoside (IPTG) and the cultures were grown for an additional 16-18 h at 16 °C. Once the TB cultures reached an OD_600_ of ∼1.0, protein expression was induced by adding 0.5 mM IPTG and the cultures were grown for an additional 16-18 h at 16 °C. Cells were pelleted at for 20 minutes at 5,000 g and 4°C. Pellets were resuspended in lysis buffer (50 mM Tris HCl pH 8.0, 200 mM NaCl, 10% Glycerol, 15 mM Imidazole, 0.1% Triton X-100, and 10 mM beta-mercaptoethanol (BME)) and the cell suspensions were placed on ice and subjected to 5 cycles of sonication at 90 A for 3 min of 1 sec on/5 sec off. The sonicated lysates were cleared by centrifugation at 15,000 rpm for 40 min at 4°C. The resulting supernatants were run over 5 mL of Ni-NTA beads that were equilibrated with lysis buffer. The column was washed with 10 column volumes (CVs) of wash buffer (50 mM Tris HCl pH 8.0, 200 mM NaCl, 25 mM Imidazole, and 10 mM BME), and then the protein of interest was eluted with 4 CVs of elution buffer (50 mM Tris HCl pH 8.0, 200 mM NaCl, 200 mM Imidazole, and 10 mM BME). The eluted protein was dialyzed overnight at 4°C using a 3.5 kDa dialysis membrane in dialysis buffer (50 mM Tris HCl pH 8.0, 200 mM NaCl, and 10 mM BME) with 3C protease added to the dialysis bag. Gel filtration chromatography was performed for each protein by loading the dialyzed sample a Superdex 75 size exclusion column (GE) in gel filtration buffer (50 mM Tris HCl pH 8.0, 200 mM NaCl, and 10 mM BME). For SCAF4, the dialyzed protein was run over Ni-NTA beads that were equilibrated with dialysis buffer prior to running size exclusion and the flowthrough was collected to load onto the size exclusion column.

### Fluorescence Polarization

All CTD peptides used in the experiment were labeled with fluorescein isothiocyanate (FITC). 40 μL reaction mixtures were set up in a black 384-well plate. CID domain proteins were titrated over a concentration range of 0-100 μM into a mixture containing 10 μM of FTIC-labeled peptide that is diluted in reaction buffer (50 mM Tris pH 8.0, 200 mM NaCl, and 10 mM BME). The plate was incubated for 1 hour at room temperature, and then fluorescence polarization values were collected using Tecan F200 plate reader. Each well was excited with vertically polarized light at 485 nm and at an emission wavelength of 535 nm. Each fluorescence polarization experiment was performed in triplicate, and GraphPad Prism v9 was used to plot and analyze the data using a one-to-one binding model to calculate *K*_d_ values.

### ChIP-Seq

HEK293T cells were seeded in 15-cm dishes and transfected with plasmids encoding YFP-RPB1_52CTD, 26CTD, or mutant constructs. After 24 hours, cells were fixed with 1% formaldehyde for 10 min at room temperature, followed by quenching with 0.125 M glycine for 5 min. Cells were sequentially lysed in 1 mL LB1 buffer (50 mM HEPES-KOH pH 7.5, 140 mM NaCl, 1 mM EDTA, 10% glycerol, 0.5% NP-40, 0.25% Triton X-100, and 1×protease and phosphatase inhibitor (1×PPI)) at 4°C for 10 min. After centrifugation and supernatant removal, the pellet was washed with 1 mL LB2 buffer (10 mM Tris-HCl pH 8.0, 200 mM NaCl, 1 mM EDTA, 0.5 mM EGTA, 1×PPI) at 4°C for 5 min. The pellet was collected by centrifugation and resuspended in 300 μL LB3 buffer (10 mM Tris-HCl pH 8.0, 100 mM NaCl, 1 mM EDTA, 0.5 mM EGTA, 0.1% Na-deoxycholate, 0.5% N-lauroylsarcosine, 1×PPI). Chromatin was sonicated to an average fragment size of approximately 200–500 bp using a Q800R3 Sonicator (30 s ON, 30 s OFF, for 20 min). For immunoprecipitation, 5 μg of GFP antibody (Proteintech, 50430-2-AP), pre-incubated with 50 μL Dynabeads Protein A (Invitrogen), was added to the chromatin samples along with 1% Triton X-100 and incubated overnight at 4°C. Beads were washed twice with low-salt wash buffer (0.1% Na-deoxycholate, 1% Triton X-100, 1 mM EDTA, 50 mM HEPES pH 7.5, 150 mM NaCl), once with high-salt wash buffer (0.1% Na-deoxycholate, 1% Triton X-100, 1 mM EDTA, 50 mM HEPES pH 7.5, 500 mM NaCl), once with LiCl wash buffer (250 mM LiCl, 0.5% NP-40, 0.5% Na-deoxycholate, 1 mM EDTA, 10 mM Tris-Cl pH 8.0), and twice with TE buffer. Chromatin was reverse crosslinked overnight at 65°C with shaking at 800 rpm in reverse crosslinking buffer (1% SDS and 0.1 M NaHCO₃). DNA was purified using phenol-chloroform extraction and resuspended in 10 mM Tris-HCl pH 8.0. Target region enrichment was confirmed by qPCR prior to deep sequencing. ChIP-seq libraries were constructed using the NEBNext Ultra II DNA Library Prep Kit and sequenced using an Illumina NovaSeq 6000 system by Novogene. Two biological replicates were performed for each condition.

### TT-Seq

Approximately 20 million cells were pulse-labeled with 1 mM 4-thiouridine (4SU, Sigma-Aldrich) for 20 min at 37 °C, then lysed in 3 mL Trizol. Total RNA was extracted, and 150 μg was used per experiment. Yeast RNA from 4TU-labeled cells was added as spike-in controls (1:100 ratio) for normalization. RNA fragmentation was achieved by adding 1 M NaOH to a final concentration of 0.2 M, followed by incubation on ice for 20 min. Fragmentation was stopped by adding 1 M Tris buffer (pH 6.8) to a final concentration of 0.5 M. RNA samples were biotinylated with MTSEA biotin-XX linker (dissolved in DMF), incubated in the dark for 40 min at room temperature, and purified using phenol/chloroform/isoamyl alcohol (25:24:1, vol/vol/vol). Biotinylated RNA was enriched using μMACS Streptavidin MicroBeads (μMACS Streptavidin Kit). Libraries were prepared using the NEBNext Ultra II Directional RNA Library Prep Kit. Two biological replicates were analyzed for each condition.

### Next-generation sequencing and data processing

The concentration and size distribution of sequencing libraries were determined by Qubit (Invitrogen) and Tapestation or Bioanalyzer (Agilent). Libraries were sequenced on Illumina NovaSeq (2x150) 6000 system by Novogene. Reads were trimmed and filtered using TrimGalore v 0.6.10 and aligned to the hg38 human genome (Ensembl) and spike-in genomes with Bowtie2 v.2.4.5 for Chip and STAR for TT-seq. PCR duplicates were removed using Picard Tools (version 3.1.1). Alignment files were converted to Bam files with Samtools (version 1.14) for downstream analysis.

### ChIP-Seq data analysis

Coverage bigWig files normalized to Input (IgG control) were generated from BAM files for each sample using the log2 ratio of mapped reads (MAPQ > 10). BigWig files from biological replicates were averaged using bigwigCompare. Gene browser tracks were visualized in IGV v2.16.2. Data matrices were generated using the computeMatrix function in Deeptools v3.5.1, from which metagene plots and heatmaps were derived. Mean signals were calculated separately for promoter region (PR, TSS to ±300 bp), gene bodies (GB, TSS+500 bp to TES) and downstream region (DS, TES+2 kb) using multiBigwigSummary (Deeptools). Box plot showing the distribution of gene-body ratios were calculated by GB / (PR + GB + DS) for yCTD and swap Pol II ChIP-seq samples were generated in R v4.4.1. ChIP-seq data generated in this study has been deposited in GEO under the BioProject accession number PRJNA1356534.

### TT-Seq data analysis

Reads mapped to exons and genes were quantified using featureCounts v2.0.1. Differential expression analysis was performed with DESeq2 v1.30.1 in R, with sizeFactors adjusted based on spike-in alignment ratios. Normalized reads were scaled accordingly, BAM files were converted to BigWig files, and replicates were averaged using bigwigCompare. Data matrices were generated using computeMatrix (Deeptools v3.5.1) to produce metagene plots. TT-seq data generated in this study has been deposited in GEO under the BioProject accession number PRJNA1356534.

### Statistics and Reproducibility

Statistical analyses were conducted using Origin Pro 9.1, RStudio v4.4.1, and GraphPad Prism 8.0. Two-tailed or one-tailed (where appropriate), independent sample t-tests were used to compare two groups. Gene-body ratio analyses were performed in R, paired Wilcoxon signed-rank tests were used to compare the distributions of gene-body ratio between yCTD and swap groups. For cell survival analyses, two-way ANOVA was employed to determine significance. Statistical source data and raw blot images are available in Source Data.

## Data Availability

Source data accompanies this manuscript. Sequencing data generated in this study have been deposited in GEO under the BioProject number: PRJNA1356534. The yCTD 26X, 52X, T4E_all, T4E_sp, and S7E_sp ChIP-seq data from a previous study (GSE252258) was used for comparison to the ChIP-seq data generated in this study.

## Acknowledgments

We thank the National Institutes of Health (R01GM104896 and R01GM125882 to YJZ, R35 GM148356 to YJZ, R01CA281106 to YJZ and CRV, R35GM142973 to RZ, and R35GM13965 to JSB) for supporting our research. We also thank L. Leon Campbell Professorship Funds for the support. Additionally, the Welch Foundation (F-1155) to JSB is gratefully acknowledged. Thanks also to Dr. Carl Wu for providing different numbers of CTD repeat constructs and Dr. Robert G. Roeder for the inducible HEK293 RPB1 degron cell line. We are thankful for helpful discussion with Bede Portz. The content is solely the responsibility of the authors and does not necessarily represent the official views of the National Institutes of Health.

## Author contributions

Q.Z. and H.H. carried out most experiments and helped with experimental design. Y. Z. and R. Z. conducted the STORM imaging experiments. M.K.V.R., J.B. and E.E. conducted mass spectrometry data analysis, A.G. helped the experimental desgin with the degron growth assay. Y.J.Z. conceived the study and experimental design. Y.J.Z., Q.Z., H.H., and R.Z. drafted the manuscript.

