## Supplemental Data for "“Zipper” Grammar of CTD Governs the Spatial Programing of the Transcription Cycle of RNA Polymerase II"

**Figure S1**

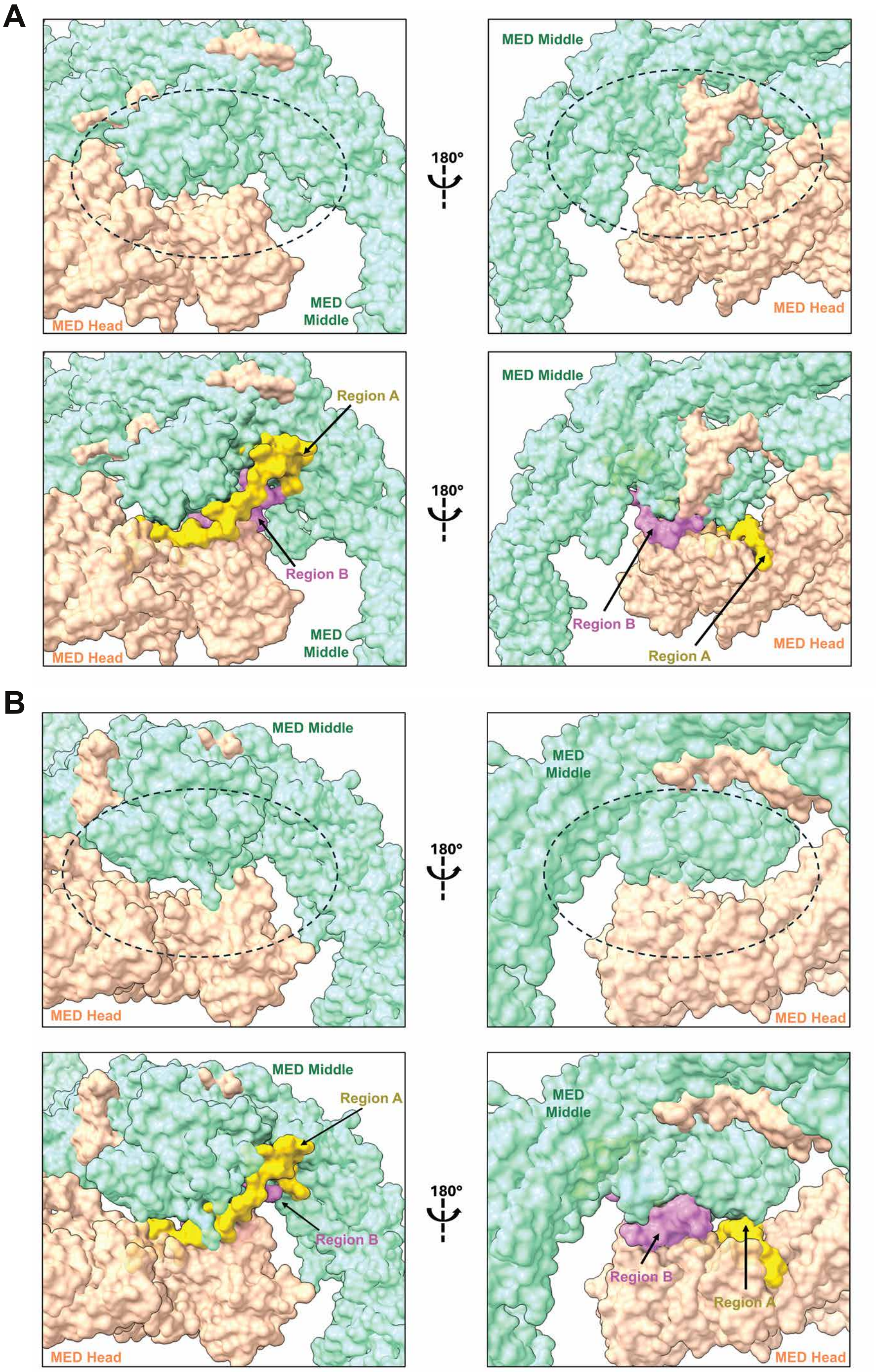


**C**

| Human MED Interface without CTD | |
|---|---|
| **Interacting Chains** | **Interface Area (Å²)** |
| n-q (Linker-Head) | 414.7 |
| m-q (Middle-Head) | 261.2 |
| d-q (Middle-Head) | 201.8 |
| g-q (Middle-Head) | 132.3 |
| q-z (Head-Middle) | 118.9 |
| m-h (Middle-Head) | 102.9 |
| d-f (Middle-Head) | 60.6 |
| **Total** | **1292.4** |

**D**

| Consensus CTD Repeats and hMED Interface | |
|---|---|
| **Interacting Chains** | **Interface Area (Å²)** |
| CTD-d (Middle) | 876.7 |
| CTD-h (Head) | 710.9 |
| CTD-f (Head) | 631.4 |
| CTD-m (Middle) | 534.3 |
| CTD-n (Linker) | 353.8 |
| CTD-q (Head) | 242.9 |
| CTD-g (Middle) | 178.5 |
| **Total** | **3528.5** |

**SI Figure 1. Structural modeling of minimal length of CTD for human PIC assembly.** (A) Front and back view of the human Mediator head and middle regions with and without the CTD bound (PDB Code: 8GXS). (B) Front and back view of the yeast Mediator head and middle regions with and without the CTD bound (PDB Code: 8CEO). (C) Summary table of the interface surface area between the human head and middle modules (calculated with PDB-ePISA). (D) Summary table of the CTD interactions with the Mediator head and middle regions in our model of the consensus CTD repeats bound to the human mediator complex.

**Figure S2**

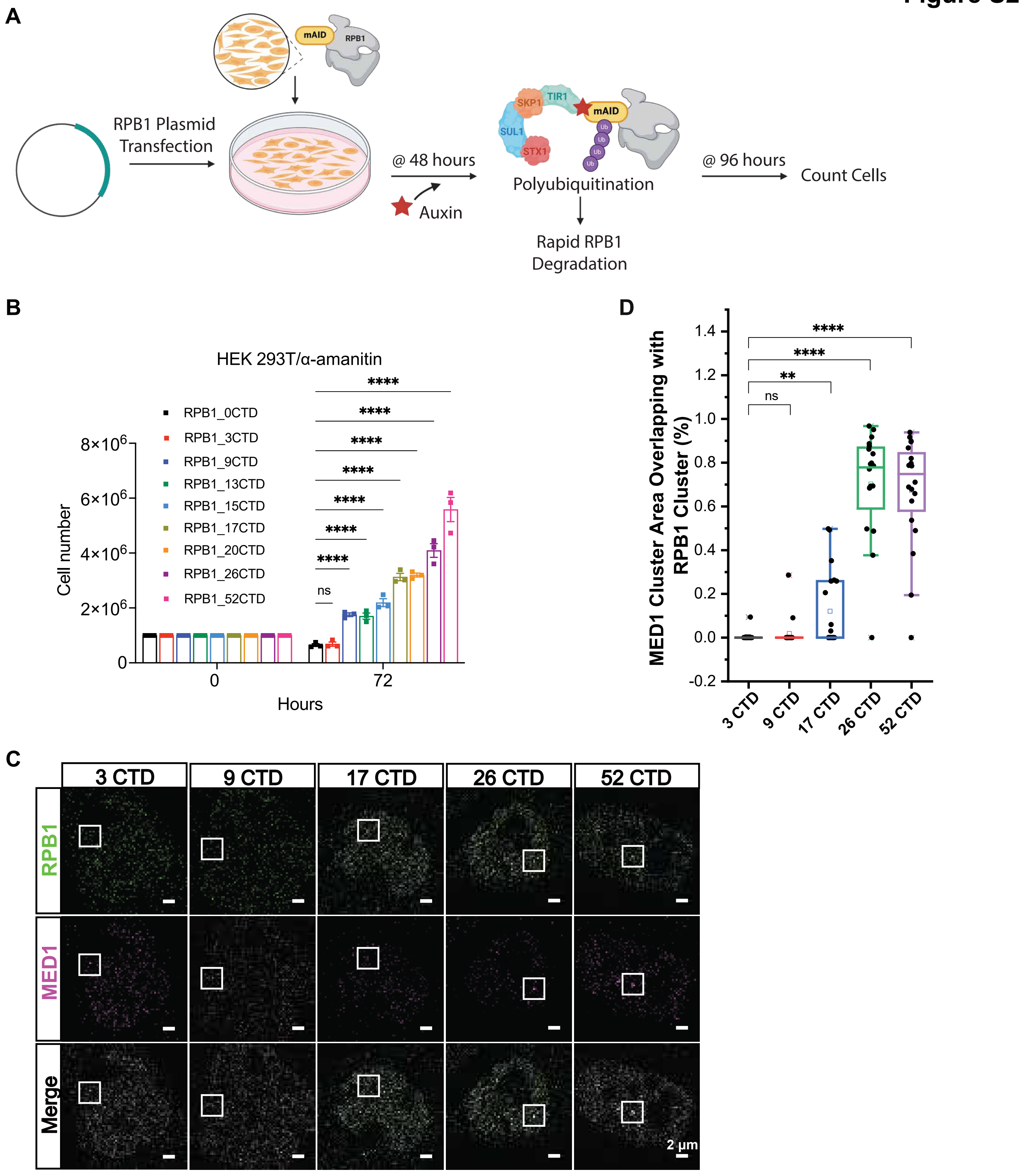


**SI Figure 2. The minimal length of CTD for cell survival and PIC assembly in cells.** (A) Standard protocol for RPB1 depletion in HEK293 RPB1 degron cells. (B) Cell density in HEK293T cells transfected wtih RPB1 that has different CTD lengths, after α-amanitin administration. n = 3 independent experiments (mean ± SEM); ****p < 0.0001, comparison was performed using two-way ANOVA with Tukey correction. (C) 3D STORM images of U2OS cells with RPB1 (Green) and MED1 (Magenta). scale bars, 2 μm. U2OS were transfected with YFP_RPB1 3 CTD, 9 CTD, 17 CTD, 26 CTD and 52 CTD. Cells were labeled with anti_GFP and anti_MED1 for two-color STORM. (D) Ratio of MED1 cluster that overlaps with RPB1. 2 μm * 2 μm area with MED1 clusters were cropped and the largest clusters for MED1 were identified for the colocalization analysis. Each data point represents a single cell. Comparison was performed using one-way ANOVA with Tukey correction.

Figure S3

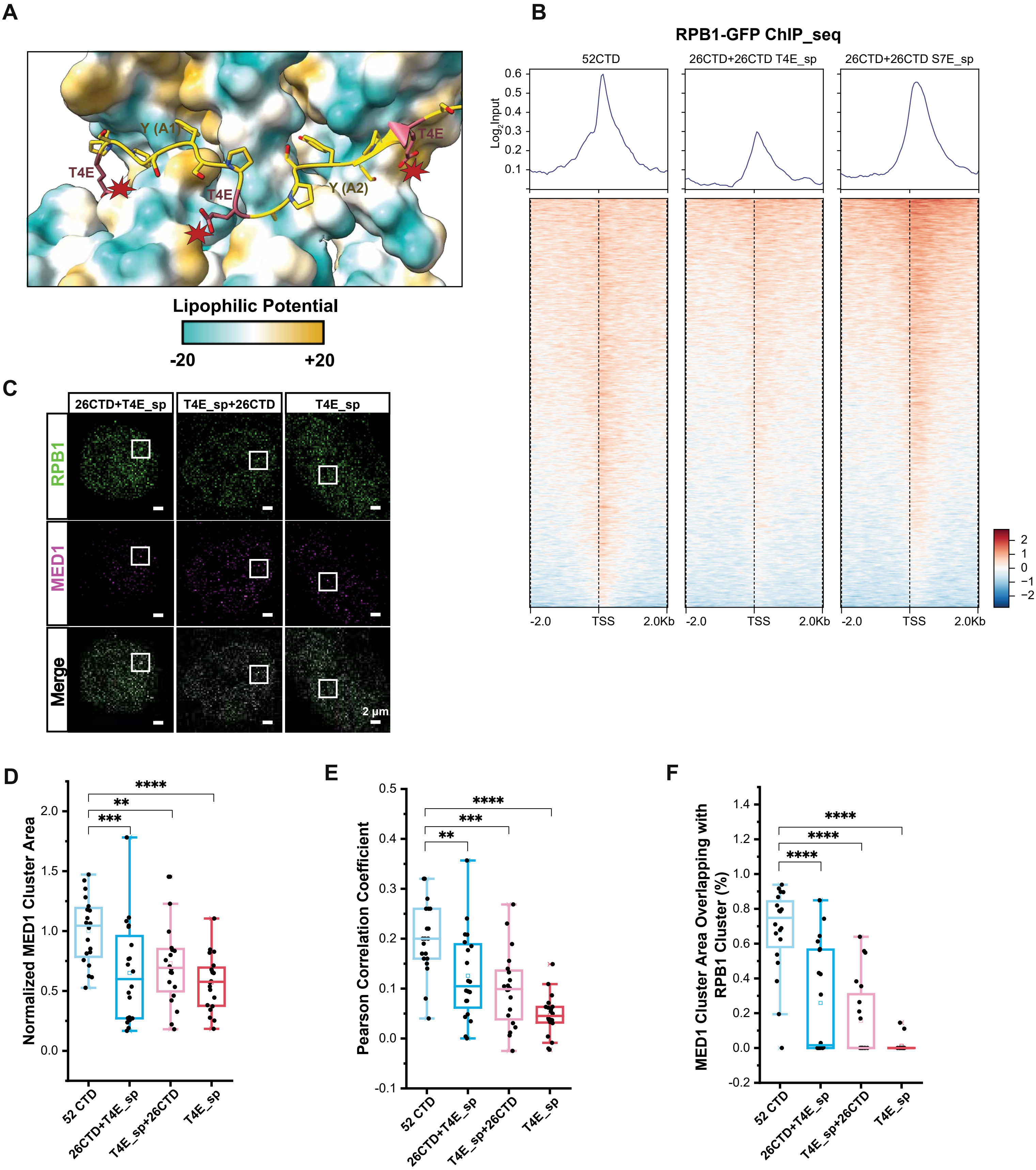


**SI Figure 3. Phosphorylation of CTD disassembles PIC and facilitates to initiation.** (A) Model of the yeast Mediator bound to a mutant T4E CTD. The Mediator (PDB Code: 8CEO) is shown as a surface representation and is colored by lipophilic potential, while the RNA Pol II CTD is shown as a cartoon. (B) Heatmaps from ChIP-seq analysis of RPB1 52CTD, 26CTD+26CTD T4E_sp and 26CTD+26CTD S7E_sp constructs. Two biological replicates (n = 2). (C) 3D STORM images of U2OS cells with RPB1 (Green) and MED1 (Magenta). scale bars, 2 µm. U2OS were transfected with YFP_RPB1 26 CTD+26CTD T4E_sp, 26CTD T4E_sp+26 CTD, and TE4_sp. Cells were labeled with anti_GFP and anti_MED1 for two-color STORM. (D) Normalized MED1 cluster area for the three constructs indicated in Fig 3H. Each data point represents a single cell. Boxplots show the mean, median and boundaries (first and third quartile); the whiskers denote the minimum and maximum excluding outliners. Comparison was performed using one-way ANOVA with Tukey correction. (E) Pearson correlation coefficients between MED1 and RPB1 for the three constructs indicated in Fig 3H. Each data point represents a single cell. Boxplots as in SI 3D. Comparison was performed using one-way ANOVA with Tukey correction. (F) Ratio of MED1 cluster that overlaps with RPB1. 2 µm * 2 µm area with MED1 clusters were cropped and the largest clusters MED1 were identified for the colocalization analysis. Each data point represents a single cell. Comparison was performed using one-way ANOVA with Tukey correction.

### Figure S4

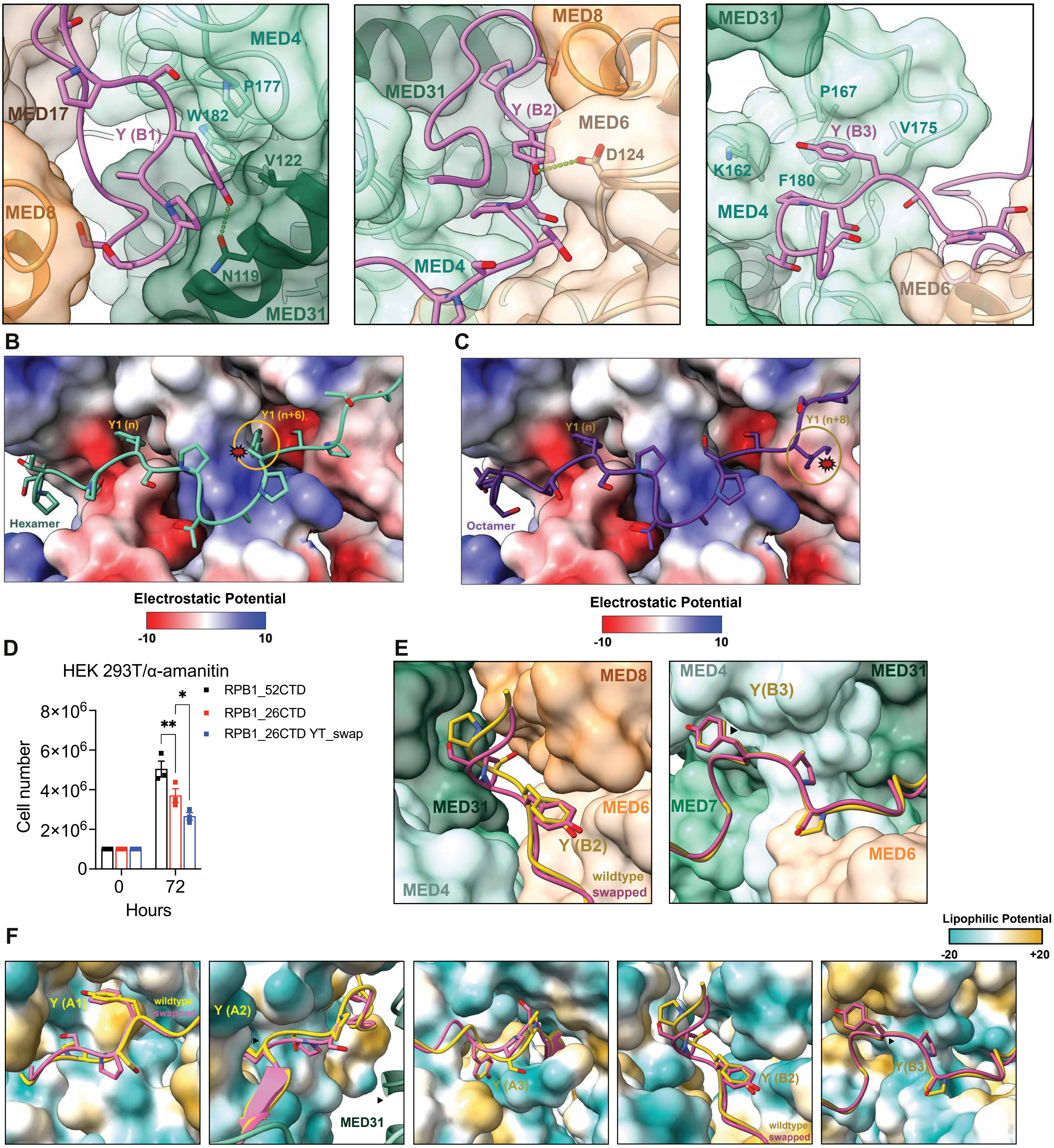


**SI Figure 4. Functional flexibility in Ser2 and Ser5 positioning within CTD heptad.** (A) Interaction between region B of the CTD with the yeast Mediator head and middle subunits (PDB Code: 8CEO). The Mediator subunits are shown as cartoons with a transparent surface representation. Relevant Mediator side chains that comprise each hydrophobic pocket or interact with the hydroxyl group of the CTD tyrosine are shown as sticks. Region B of the CTD is shown in pink. (B) Model of the yeast Mediator bound to a mutant hexamer CTD. The Mediator is shown as a surface representation and is colored by electrostatic potential, while the Pol II CTD is shown as a cartoon. (C) Model of the yeast Mediator bound to a mutant octamer CTD. The Mediator is shown as a surface representation and is colored by electrostatic potential, while the Pol II CTD is shown as a cartoon. (D) Cell density of HEK293T cells transfected with RPB1 52CTD, 26CTD and 26CTD YT_swap after α-amanitin administration. n = 3 independent experiments (mean ± SEM); ****p < 0.0001, comparison was performed using two-way ANOVA with Tukey correction. (E) Region B of the wild type CTD (yellow) or a model of the swapped CTD (pink) bound to the yeast Mediator. Mediator head subunits are colored varying shades of orange, while Mediator middle subunits are colored varying shades of green. (F) Regions A and B of the wild type CTD (yellow) or a model of the swapped CTD (pink) bound to the yeast Mediator. Mediator head subunits are colored by lipophilic potential.

### Figure S5

**A**

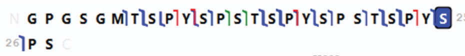


**2**

| Identified Ions | Theoretical Mass | Observed Mass | Mass Difference (ppm) |
|---|---|---|---|
| A+11 | 994.4304 | 994.4266 | -3.83137 |
| A+12 | 1091.483 | 1091.483 | -0.0834 |
| A+13 | 1178.515 | 1178.507 | -7.12934 |
| A+16 | 1463.648 | 1463.648 | 0.345006 |
| A+17 | 1626.711 | 1626.707 | -2.54319 |
| A+6 | 459.2026 | 459.1995 | -6.69864 |
| A10 | 906.3905 | 906.394 | 3.80741 |
| A15 | 1365.587 | 1365.587 | -0.06517 |
| A22 | 2084.9 | 2084.902 | 1.284474 |
| A24 | 2345.016 | 2345.019 | 1.171847 |
| B10 | 934.3855 | 934.3868 | 1.453362 |
| B11 | 1021.417 | 1021.419 | 1.299175 |
| B14 | 1306.55 | 1306.551 | 0.813593 |
| B15 | 1393.582 | 1393.585 | 1.961851 |
| B16 | 1490.635 | 1490.649 | 9.693857 |
| B17 | 1653.698 | 1653.7 | 0.907663 |
| B18 | 1740.73 | 1740.733 | 1.479264 |
| B20 | 1924.815 | 1924.807 | -3.97649 |
| B21 | 2025.863 | 2025.863 | 0.248289 |
| B22 | 2112.895 | 2112.898 | 1.691518 |
| B24 | 2373.011 | 2373.015 | 1.983556 |
| B25 | 2540.009 | 2540.014 | 1.762198 |
| B6 | 486.1897 | 486.19 | 0.730168 |
| B7 | 587.2373 | 587.2381 | 1.212457 |
| C10 | 951.4118 | 951.4175 | 6.049957 |
| C16 | 1507.661 | 1507.663 | 1.070532 |
| C23 | 2226.974 | 2226.973 | -0.47823 |
| C9 | 788.3484 | 788.3483 | -0.13065 |
| Y10 | 1088.406 | 1088.407 | 0.923368 |
| Y12 | 1348.522 | 1348.53 | 5.950958 |
| Y13 | 1435.554 | 1435.555 | 0.13305 |
| Y17 | 1807.719 | 1807.722 | 1.523467 |
| Y19 | 2067.835 | 2067.84 | 2.624 |
| Y20 | 2154.867 | 2154.866 | -0.50954 |
| Y3 | 369.0937 | 369.094 | 0.823639 |
| Y5 | 629.2098 | 629.2105 | 1.056881 |
| Y6 | 716.2418 | 716.2425 | 0.973135 |

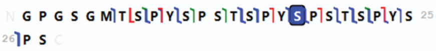


**3**

| Identified Ions | Theoretical Mass | Observed Mass | Mass Difference (ppm) |
|---|---|---|---|
| A+11 | 994.4304 | 994.4225 | -7.96338 |
| A+17 | 1626.711 | 1626.708 | -1.76063 |
| A+22 | 2165.874 | 2165.867 | -3.26706 |
| A+6 | 459.2026 | 459.2003 | -4.95649 |
| A+8 | 647.2823 | 647.2784 | -6.07159 |
| A+9 | 744.335 | 744.334 | -1.41742 |
| A10 | 906.3905 | 906.3927 | 2.335638 |
| A13 | 1177.507 | 1177.501 | -5.54646 |
| A15 | 1365.587 | 1365.589 | 1.04424 |
| A25 | 2512.014 | 2512.012 | -0.91441 |
| B10 | 934.3855 | 934.387 | 1.627808 |
| B14 | 1306.55 | 1306.552 | 1.679997 |
| B15 | 1393.582 | 1393.584 | 1.79035 |
| B17 | 1653.698 | 1653.701 | 1.675034 |
| B18 | 1820.696 | 1820.706 | 5.389696 |
| B20 | 2004.781 | 2004.792 | 5.511823 |
| B21 | 2105.829 | 2105.828 | -0.20894 |
| B24 | 2452.977 | 2452.978 | 0.481456 |
| B25 | 2540.009 | 2540.014 | 1.989363 |
| B6 | 486.1897 | 486.19 | 0.742509 |
| B9 | 771.3221 | 771.3283 | 8.022588 |
| C16 | 1507.661 | 1507.663 | 0.98298 |
| C19 | 1934.776 | 1934.763 | -6.28548 |
| C23 | 2306.94 | 2306.941 | 0.577822 |
| C9 | 788.3484 | 788.3488 | 0.502316 |
| Y10 | 1088.406 | 1088.411 | 4.456056 |
| Y13 | 1435.554 | 1435.556 | 1.402942 |
| Y17 | 1807.719 | 1807.723 | 2.110948 |
| Y19 | 2067.835 | 2067.835 | 0.139276 |
| Y4 | 452.1907 | 452.1923 | 3.410066 |
| Y5 | 549.2435 | 549.2442 | 1.332742 |
| Y6 | 636.2755 | 636.2763 | 1.277748 |
| Y7 | 737.3232 | 737.3222 | -1.30879 |
| Y9 | 921.408 | 921.4088 | 0.894284 |
| Z20 | 2138.848 | 2138.835 | -6.34781 |

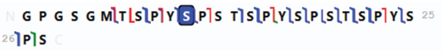


**4**

| Identified Ions | Theoretical Mass | Observed Mass | Mass Difference (ppm) |
|---|---|---|---|
| A+15 | 1446.561 | 1446.562 | 0.736898 |
| A+25 | 2513.022 | 2513.018 | -1.65181 |
| A+8 | 647.2823 | 647.2784 | -5.931 |
| A+9 | 744.335 | 744.3332 | -2.4707 |
| A10 | 906.3905 | 906.3923 | 1.925219 |
| A11 | 1073.389 | 1073.396 | 6.565188 |
| A22 | 2164.866 | 2164.875 | 4.159149 |
| A26 | 2609.067 | 2609.079 | 4.63499 |
| B10 | 934.3855 | 934.3866 | 1.195438 |
| B11 | 1101.384 | 1101.39 | 5.891679 |
| B14 | 1386.516 | 1386.518 | 1.436694 |
| B15 | 1473.548 | 1473.551 | 1.915784 |
| B17 | 1733.664 | 1733.667 | 1.511827 |
| B18 | 1820.696 | 1820.699 | 1.642778 |
| B20 | 2004.781 | 2004.797 | 8.024816 |
| B21 | 2105.829 | 2105.829 | 0.16573 |
| B22 | 2192.861 | 2192.865 | 1.95726 |
| B24 | 2452.977 | 2452.979 | 0.673467 |
| B25 | 2540.009 | 2540.014 | 1.789757 |
| B6 | 486.1897 | 486.1906 | 1.86347 |
| B8 | 674.2694 | 674.27 | 0.9032 |
| B9 | 771.3221 | 771.3164 | -7.44566 |
| C12 | 1215.463 | 1215.46 | -2.04037 |
| C16 | 1587.627 | 1587.63 | 1.546962 |
| C23 | 2306.94 | 2306.948 | 3.582234 |
| C6 | 503.216 | 503.2164 | 0.965788 |
| C9 | 788.3484 | 788.3483 | -0.15095 |
| X+11 | 1198.49 | 1198.486 | -3.9283 |
| Y10 | 1008.44 | 1008.441 | 0.753639 |
| Y12 | 1268.556 | 1268.558 | 1.381886 |
| Y16 | 1640.721 | 1640.721 | 0.116412 |
| Y17 | 1807.719 | 1807.722 | 1.689422 |
| Y19 | 2067.835 | 2067.836 | 0.365116 |
| Y-21 | 2254.907 | 2254.889 | -7.79499 |
| Y3 | 289.1274 | 289.1276 | 0.864671 |
| Y5 | 549.2435 | 549.2441 | 1.172522 |
| Y5 | 549.2435 | 549.2441 | 1.172522 |
| Y6 | 636.2755 | 636.276 | 0.777965 |
| Y7 | 737.3232 | 737.3287 | 7.466197 |
| Y8 | 824.3552 | 824.3545 | -0.82489 |
| Y9 | 921.408 | 921.4089 | 1.02669 |
| Y-9 | 920.4001 | 920.4086 | 9.199295 |
| Z20 | 2138.848 | 2138.849 | 0.225355 |

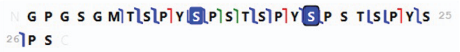


**5**

| Identified Ions | Theoretical Mass | Observed Mass | Mass Difference (ppm) |
|---|---|---|---|
| A+12 | 1171.449 | 1171.446 | -2.759 |
| A13 | 1257.474 | 1257.466 | -6.46216 |
| B11 | 1101.384 | 1101.383 | -0.73635 |
| B14 | 1386.516 | 1386.517 | 0.571216 |
| B15 | 1473.548 | 1473.549 | 0.716638 |
| B17 | 1733.664 | 1733.667 | 1.500867 |
| B24 | 2532.943 | 2532.945 | 0.647863 |
| B25 | 2619.975 | 2619.98 | 1.590473 |
| B6 | 486.1897 | 486.1882 | -2.93918 |
| B7 | 587.2373 | 587.2377 | 0.602823 |
| B8 | 674.2694 | 674.2693 | -0.10382 |
| C16 | 1587.627 | 1587.63 | 1.520508 |
| C23 | 2386.906 | 2386.909 | 1.324727 |
| C9 | 788.3484 | 788.3487 | 0.394496 |
| Y10 | 1088.406 | 1088.413 | 6.153952 |
| Y13 | 1435.554 | 1435.554 | -0.39776 |
| Y16 | 1720.687 | 1720.687 | 0.021503 |
| Y17 | 1887.685 | 1887.687 | 0.793035 |
| Y19 | 2147.801 | 2147.802 | 0.221156 |
| Y20 | 2234.833 | 2234.84 | 3.044522 |
| Y3 | 289.1274 | 289.1275 | 0.408125 |
| Y3 | 289.1274 | 289.1275 | 0.408125 |
| Y5 | 549.2435 | 549.2436 | 0.18571 |
| Y6 | 636.2755 | 636.2754 | -0.23418 |
| Y-6 | 635.2677 | 635.2653 | -3.76529 |
| Y9 | 921.408 | 921.4138 | 6.310994 |

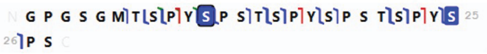


**6**

| Identified Ions | Theoretical Mass | Observed Mass | Mass Difference (ppm) |
|---|---|---|---|
| A+8 | 647.2823 | 647.2762 | -9.44106 |
| A+9 | 744.335 | 744.3337 | -1.82046 |
| A10 | 906.3905 | 906.3918 | 1.382406 |
| B11 | 1101.384 | 1101.389 | 4.526124 |
| B13 | 1285.469 | 1285.479 | 8.234351 |
| B14 | 1386.516 | 1386.52 | 2.7277 |
| B15 | 1473.548 | 1473.55 | 0.900547 |
| B17 | 1733.664 | 1733.668 | 1.947897 |
| B18 | 1820.696 | 1820.699 | 1.160545 |
| B21 | 2105.829 | 2105.832 | 1.309223 |
| B22 | 2192.861 | 2192.871 | 4.670155 |
| B24 | 2452.977 | 2452.983 | 2.353874 |
| B25 | 2619.975 | 2619.98 | 1.795437 |
| B6 | 486.1897 | 486.1897 | 0.168658 |
| C16 | 1587.627 | 1587.63 | 1.577196 |
| C23 | 2306.94 | 2306.948 | 3.331686 |
| C9 | 788.3484 | 788.3489 | 0.613942 |
| Y10 | 1088.406 | 1088.409 | 2.719573 |
| Y13 | 1435.554 | 1435.558 | 2.537695 |
| Y-16 | 1719.679 | 1719.677 | -1.01936 |
| Y17 | 1887.685 | 1887.686 | 0.153627 |
| Y19 | 2147.801 | 2147.806 | 2.31865 |
| Y20 | 2234.833 | 2234.834 | 0.200015 |
| Y3 | 369.0937 | 369.0941 | 0.967234 |
| Y6 | 716.2418 | 716.2429 | 1.486928 |

**B**

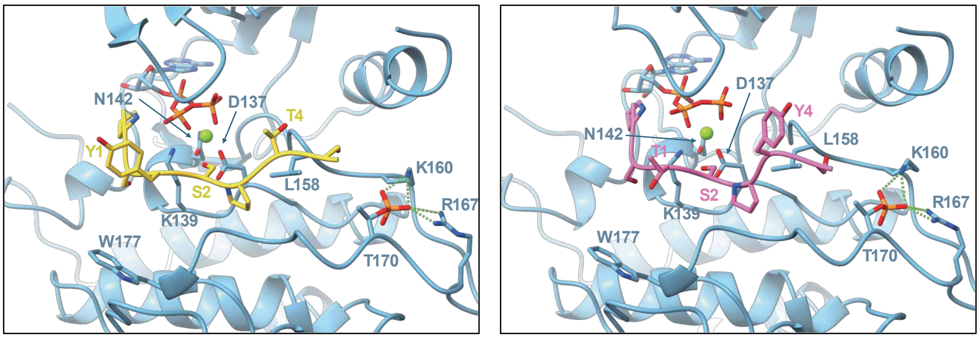


CDK7 with pSer2 in active site (WT CTD)

CDK7 with pSer2 in active site (swapped CTD)

**SI Figure 5. Ser2/Ser5 Positional Swap Does Not Impair CTD Phosphorylation or PIC Disassembly.** (A) Lists of fragment ions for identification of mono-phosphorylated peptides (m/z 915.38) and di-phosphorylated (m/z 942.04) analyzed in Figure 5C by UVPD-MS. In each case, the 3+ charge state was selected, and UVPD was performed using 2 laser pulses (1.5 mJ per pulse). The identified sites of phosphorylation are shaded in blue in the sequence map above each table. Fragment ions are named as Tn where T is the type of ion (A = a, B = b, C = c, X = x, Y = y, Z = z, for which A,B, and C originate from the N-terminus of the peptide and X,Y and Z originate from the C-terminus of the peptide), the subscript indicates the number of amino acids contained in the fragment ion, and a plus or minus sign in the subscript designates whether the fragment ion contains one extra hydrogen atom or lacks one hydrogen atom. (B) Model of CDK7 bound to the RNA Pol II CTD (yellow, left) and model of CDK7 bound to the swapped RNA Pol II CTD (pink, right). Ser2 was positioned for phosphorylation within the CDK7 active site for both the wild type and swapped CTD.

### Figure S6

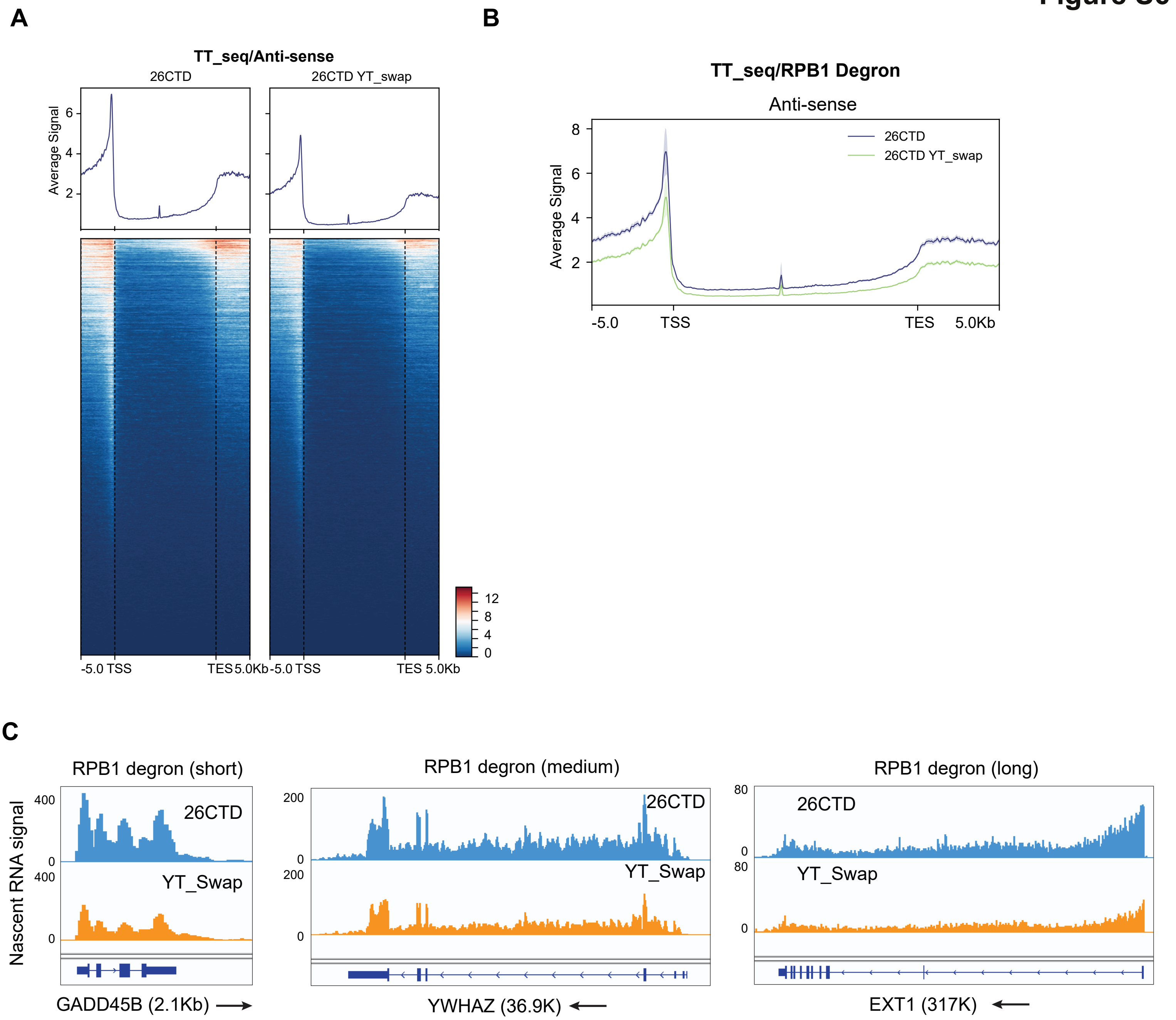


**SI Figure 6. Ser2/Ser5 swapping affects transcription elongation and 3' end processing.** (A) Average antisense signal of heatmaps in 26CTD and YT_swap CTD for TTseq, 2 replicates for each group. (B) Metagene plots for antisense profile of 26CTD and YT_swap CTD group. (C) Browser tracks of TTseq data at short (GADD45B), medium (YWHAZ), and long genes (EXT1).
